# Flexible Sensory-Motor Mapping Rules Manifest in Correlated Variability of Stimulus and Action Codes Across the Brain

**DOI:** 10.1101/2022.03.10.483758

**Authors:** R.L. van den Brink, K. Hagena, N. Wilming, P.R. Murphy, C. Büchel, T.H. Donner

## Abstract

Humans and non-human primates can flexibly switch between different arbitrary mappings from sensation to action to solve a cognitive task. It has remained unknown how the brain implements such flexible sensory-motor mapping rules. Here, we uncovered a dynamic reconfiguration of task-specific correlated variability between sensory and motor brain regions. Human participants switched between two rules for reporting visual orientation judgments during fMRI recordings. Rule switches were either signaled explicitly or inferred by the participants from ambiguous cues. We used behavioral modeling to reconstruct the time course of their belief about the active rule. In both contexts, the patterns of correlations between ongoing fluctuations in stimulus- and action-selective activity across visual and action-related brain regions tracked participants’ belief about the active rule. The rule-specific correlation patterns broke down around the time of behavioral errors. We conclude that internal beliefs about task state are instantiated in brain-wide, selective patterns of correlated variability.

## Introduction

Perceptual decisions transform an internal representation of the state of the environment into motor output in a flexible, context-dependent manner (Mante et al., 2013; Shadlen and Kiani, 2013). Substantial progress has been made in uncovering the neural bases of such transformations during tasks in which the required mapping from sensation to action was fixed (Bogacz et al., 2006; Gold and Shadlen, 2007; Donner et al., 2009; Hanks et al., 2015; Wilming et al., 2020; Murphy et al., 2021). Such task-relevant sensory-motor mappings often entail an arbitrary association of specific features of the sensory input with specific features of the motor output (Sakai, 2008; Shadlen and Kiani, 2013).

Consider the mapping rule that governs a canonical perceptual choice task: visual orientation discrimination (Figure 1). The task rule may require reporting a vertical orientation of a visual grating with by button press with the left hand and horizontal with right hand (Figure 1a, Mapping rule 1), or the converse (Figure 1a, Mapping rule 2). Primate behavior rapidly tracks switches between such distinct sensory-motor mapping rules (Gold and Shadlen, 2003; Tsetsos et al., 2015; Purcell and Kiani, 2016). To accomplish this, the brain must flexibly configure task-specific networks that link certain sensory features with certain motor actions (Shadlen and Kiani, 2013). In our example task, signals from vertical and horizontal orientation columns in primary visual cortex (V1) (Bonhoeffer and Grinvald, 1991) need to be routed to distinct neural populations in primary motor cortex (M1) that control left- or right-hand movements, respectively (Yousry et al., 1997) under each rule.

**Figure 1.**
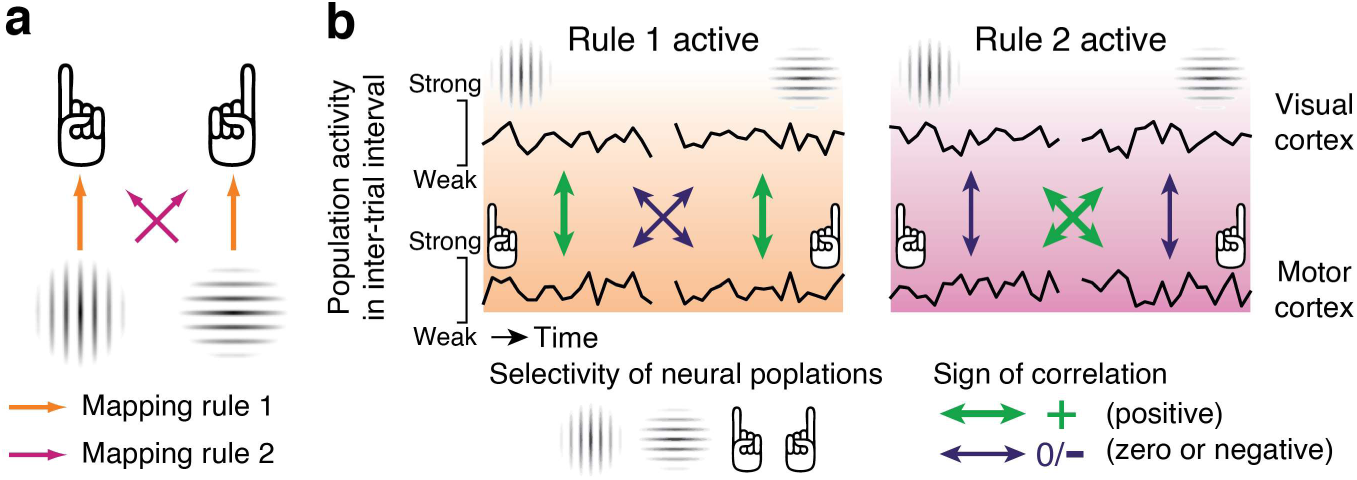
Sensory-motor mapping rules, task-relevant functional networks, and correlated neural variability. **a.** Example sensory-motor mapping rule in a visual orientation discrimination task. Colored arrows indicate the mappings from sensory input to motor output dictated by the two rules. **b.** Predicted patterns of correlated variability between orientation-selective populations of visual cortical neurons (top) and action-selective populations of motor cortical neurons (bottom). Stimulus- and action-coding evoked responses during task are bound to correlate in a rule-specific fashion when subjects follow the rule. Our hypothesis focusses on ongoing (i.e., not stimulus- or task-evoked) fluctuations in activity, over and above the evoked responses. These fluctuations are shown for two intervals between the trials of the orientation discrimination task, one interval for each rule. Green and purple arrows; the pattern of intrinsic correlations between stimulus- and action-selective populations flips between rules.

How this dynamic reconfiguration of task-specific networks is accomplished has remained elusive. A large body of work has quantified the costs of switching between different ‘task sets’ (Sakai, 2008), assessed behaviorally or in terms of the recruitment of brain regions involved in cognitive control (Monsell, 2003). Another line of prior research has studied the encoding of rule information within individual brain regions (Wallis et al., 2001; Stoet and Snyder, 2004; Haynes et al., 2007; Sakai, 2008; Bode and Haynes, 2009; Bennur and Gold, 2011; Zhang et al., 2013; Cole et al., 2016). Here, we instead asked whether and how the rule may shape the patterns of correlations of ongoing (not stimulus- or task-evoked) activity between sensory and action-related regions. Such intrinsic correlations of brain activity, often referred to as ‘functional connectivity’, delineate functional networks in the brain (Gerstein and Perkel, 1969; Büchel and Friston, 2000; Vincent et al., 2007). Rule-dependent changes in intrinsic correlations may result from dynamic modifications of task-relevant sensory-motor cortical connections through short-term plasticity mechanisms (Fusi et al., 2007). A (not mutually exclusive) alternative is that such correlations result from selective common input to sensory and motor brain regions, for example from rule-encoding neural populations in prefrontal cortex (Miller and Cohen, 2001).

Whatever their source, we reasoned that these correlations should be expressed between the specific neural populations that encode the task-relevant sensory features (e.g., horizontal or vertical) and action features (e.g., left- or right-hand button press; Figure 1b). In other words, rule-specific correlations should couple fine-grained feature-coding activity patterns across different brain regions, rather than these regions’ overall activity as commonly assessed in fMRI studies of functional connectivity (Büchel and Friston, 2000; Vincent et al., 2007). Furthermore, the correlations should be expressed in ongoing activity, after accounting for evoked responses and even during the intervals between the trials of the primary task (Figure 1b). In our example task, maintenance of mapping rule 1 should be reflected in co-fluctuations between vertical-preferring neural populations in visual cortex, and left-hand movement-encoding neurons in motor cortex. Horizontal-preferring visual cortical populations should similarly co-fluctuate with right-hand movement-encoding motor cortical neurons. Conversely, the other two pairs of neural populations should be decoupled (or anti-correlated; Figure 1b, orange). When the rule then switches to rule 2, this selective pattern of correlated variability should flip (Figure 1b, pink).

We developed an approach to test this idea in human fMRI recordings. We asked participants to perform a visual orientation discrimination task under varying sensory-motor mapping rules and analyzed the fine structure of intrinsic correlations between stimulus- and action-selective patterns of fMRI activity expressed in different brain areas. In another version of the task, we let rules switch in a hidden and unpredictable fashion, which required participants to use a rapidly evolving, adaptive inference process for tracking the changing rules uncertainty, giving rise to frequent errors in their rule-guided behavior. We reasoned that the intrinsic fMRI signal correlations should then mirror the evolution of participants’ hidden, graded beliefs belief about the active rule. Behavioral modeling enabled us to reconstruct these belief states at any given moment.

We found that the patterns of intrinsic correlations between stimulus- and action-selective population activity reliably tracked participants’ belief states, but selectively broke down around the time of behavioral errors. This links the structure of correlated variability to cognitive computation and establishes its significance for flexible sensory-motor decisions.

## Results

We alternated the stimulus-response mapping rule (SR-rule) of a simple perceptual choice task (Figure 1a). In one set of fMRI runs (see Methods), the SR-rule for the upcoming trials was explicitly instructed by visual cues (SR-rule cue), which defined mini-blocks of two trials each (Figure 2a-b). We call this setting ‘instructed rule’. Each trial of the primary task entailed the presentation of a large, full-contrast, horizontal or vertical visual grating stimulus, followed by the participants’ report of their orientation judgment. Grating stimuli were followed by long and variable inter-trial intervals (Figure 2a).

**Figure 2.**
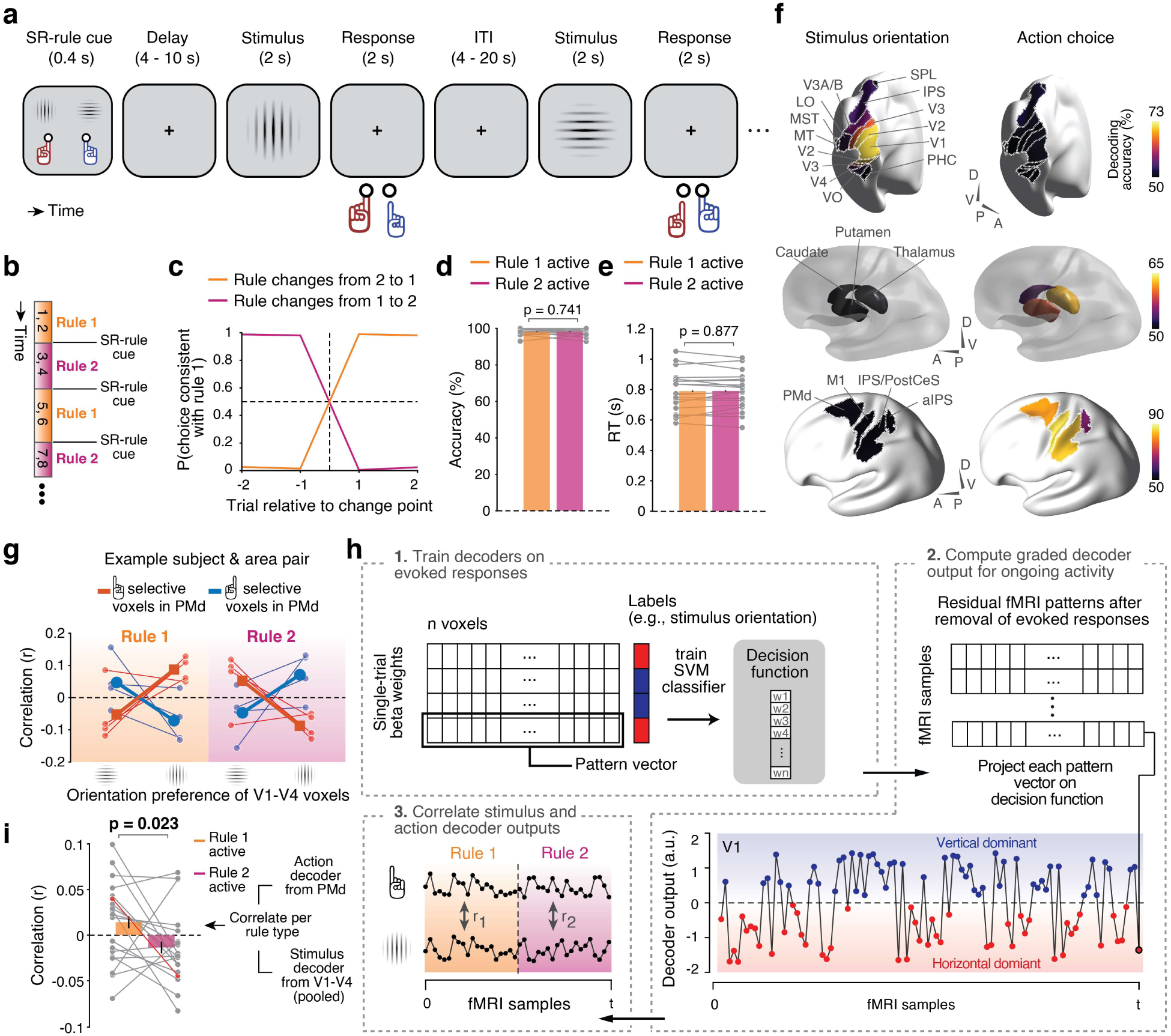
Instructed rule task and assessment of feature-selective correlated variability. **a, b.** Instructed rule task. **a.** Example sequence of events during a mini-block consisting of two trials. The SR-rule cue instructs the rule for the following mini-block. After a variable delay, two trials of the primary task are presented, each followed by a variable inter-trial interval (ITI). See main text for details. **b.** Mini-block structure of the task. Numbers in the mini-blocks indicate trials. **c.** Group average time course of choice fractions locked to instructed rule changes. Shaded area, between-participant SEM. For many data points, this area is smaller than the line for the mean. **d, e.** Overall behavioral accuracy (d) and mean RT (e), separately for the two rules. Gray dots, individual participants; large red/blue dots, group-averages. **f.** Classification of stimulus orientation and action by decoders trained and tested on evoked responses during primary task trials. Decoding performance is evaluated for 19 pre-selected cortical and subcortical regions (pooled across hemispheres). **g.** Selective co-fluctuation pattern for feature-selective voxel groups from example participant and region pair (V1-V4 and PMd) during segments of time in which rule 1 (orange) or rule 2 (pink) is active. This and all subsequent analyses of fMRI correlations account for hemodynamic lag (Materials and Methods). Thin lines, individual task runs (N = 4); thick lines, average. **h.** General analysis approach. See main text and Methods for details. **i.** Correlations between stimulus and action decoder outputs for the same area pair as in panel g, evaluated during the two rules. Thin lines, individual participants (N = 19); red line, example subject from panel g; bars, group average; error bars, within-participant SEM. All *p*-values based on permutation tests; boldface, *p* < 0.05 (Table S1).

Participants were able to switch rapidly and consistently between the rules (Figure 2c), reaching choice fractions near-fully consistent with the active rule on the first trial after the switch. All participants’ overall performance was close to ceiling (Figure 2d), with no differences between the two rules in accuracy (Figure 2d), RT (Figure 2e), or fixational eye movements (Figure S1). High overall accuracy was expected because the task entailed no uncertainty about the active rule nor about the primary perceptual judgment (maximal contrast and orientation difference between discriminanda). Equal performance for both rules was expected because the rules were fully symmetric, without any obvious benefit of using one rather than the other rule (as opposed to, e.g., the so-called Simon effect (Simon and Wolf, 1963), see Discussion). This setting yielded clear expectations for the dynamics of putative changes in patterns of correlated variability during rule switching (Figure 1b).

### Correlated variability of stimulus and action codes reflects instructed rule

We reasoned that if the sensory-motor mapping rule sculpts the sensory-motor network that implements the primary choice task, the active rule should be reflected in co-fluctuations between stimulus- and action-selective populations of neurons in sensory and motor brain regions, respectively (Figure 1b). To test this idea, we analyzed correlations between fMRI signal fluctuations measured in feature-specific subsets of voxels in these different brain regions.

To illustrate the rationale of our approach, we first focus on two sets of regions implicated in visual orientation perception and action planning or execution, respectively: early visual cortex (V1-V4) (Reynolds et al., 2000; Haynes and Rees, 2005a; Kamitani and Tong, 2005) and dorsal premotor cortex (PMd; Figure 2f) (Picard and Strick, 2001; Cisek and Kalaska, 2005; Murphy et al., 2021; Peixoto et al., 2021). Here, and for illustration only, we selected subgroups of voxels that responded preferentially to vertical or horizontal gratings (for V1-V4) or to left- or right-hand choices (PMd). For both areas, we then removed the evoked responses of each individual voxel (using linear regression), in order to isolate multi-voxel patterns of ongoing fluctuations of activity in the residual (Arieli et al., 1996; Kenet et al., 2003; Fox et al., 2006). We then correlated the residuals between all four pairs of voxel sub-groups (Methods). In the example participant in Figure 2g, the resulting patterns of correlations corresponded to our qualitative prediction from Figure 1b: for rule 1, we found positive correlations for vertical and left-hand coding voxels and for horizontal and right-hand coding voxels, and negative correlations for the other two pairs (Figure 2g, orange), and this pattern flipped for rule 2 (Figure 2g, pink).

To quantify these intrinsic correlation patterns in a more compact fashion across the group of participants and all pairs of areas, we trained stimulus- and action decoders on the response patters evoked during trials of the primary task (Figure 2h, box 1; see Methods for voxel selection and training procedure). As expected (e.g. Haynes and Rees, 2005b; Kamitani and Tong, 2005; de Gee et al., 2017), stimulus orientation was robustly decodable from patterns of evoked responses in several visual cortical areas, and action choice was robustly decodable from evoked responses in areas of the cortical motor system (anterior parietal cortex, PMd, and M1; Figures 2f and S2). We also found above-chance action (but not stimulus) decoding from subcortical regions (thalamus, caudate, and putamen; Figures 2f and S2). We then applied these decoders to the ongoing (i.e., residual) fluctuations of activity patterns throughout the recording (Methods and Figure 2h, box 2). These residuals neither exhibited an overall (voxel-average) evoked response, nor any stimulus- or action-selective multi-voxel activity patterns that were evoked by (i.e., systematically time-locked to) the trials of the primary task (Figure S3). The resulting time courses of decoder outputs compactly summarized the fluctuations of the feature-selective activity component within each region. For example, in V1, the time course reflected the extent to which patterns of ongoing population activity (Kenet et al., 2003) tended towards vertical (positive sign) or horizontal (negative) orientations (Figure 2h, box 2). We finally correlated stimulus and action decoder outputs, separately for intervals corresponding to the two rules (Figure 2h, box 3).

Given the way we constructed the decoders for stimulus (positive for vertical and negative prediction for horizontal) and action (positive output for left hand and negative output for right hand; Figure 2h), our hypothesis (Figure 1b) predicted that the correlation of decoder outputs in the ongoing activity should be positive when rule 1 was active and negative when rule 2 was active (Figure 2h, box 3). This is precisely what we found for the output of the V1-V4 stimulus decoder and the PMd action decoder, both in our example participant (see Figure 2i, red line) as well as in the whole group (Figure 2i, gray lines and colored bars). Thus, the co-fluctuations of feature-selective patterns of ongoing activity in early visual cortex and PMd flipped sign depending on the active rule.

This sign flip between rules was evident for most pairs of stimulus- and action-encoding regions (Figure 3): The correlation between stimulus and action decoder outputs (gray rectangles in Figure 3a-c), was positive during rule 1 for most region pairs (Figure 3a) and negative under rule 2 for most region pairs (Figure 3b). This yielded a correlation difference that was consistent across most pairs of stimulus-encoding and action-encoding regions (Figure 3c, cells inside gray rectangle). Correspondingly, collapsing the correlations across all stimulus-action decoder pairs (gray rectangle in Figure 3c) yielded a positive correlation for rule 1 and negative correlation for rule 2, with a clear difference between the two (Figure 3e).

**Figure 3.**
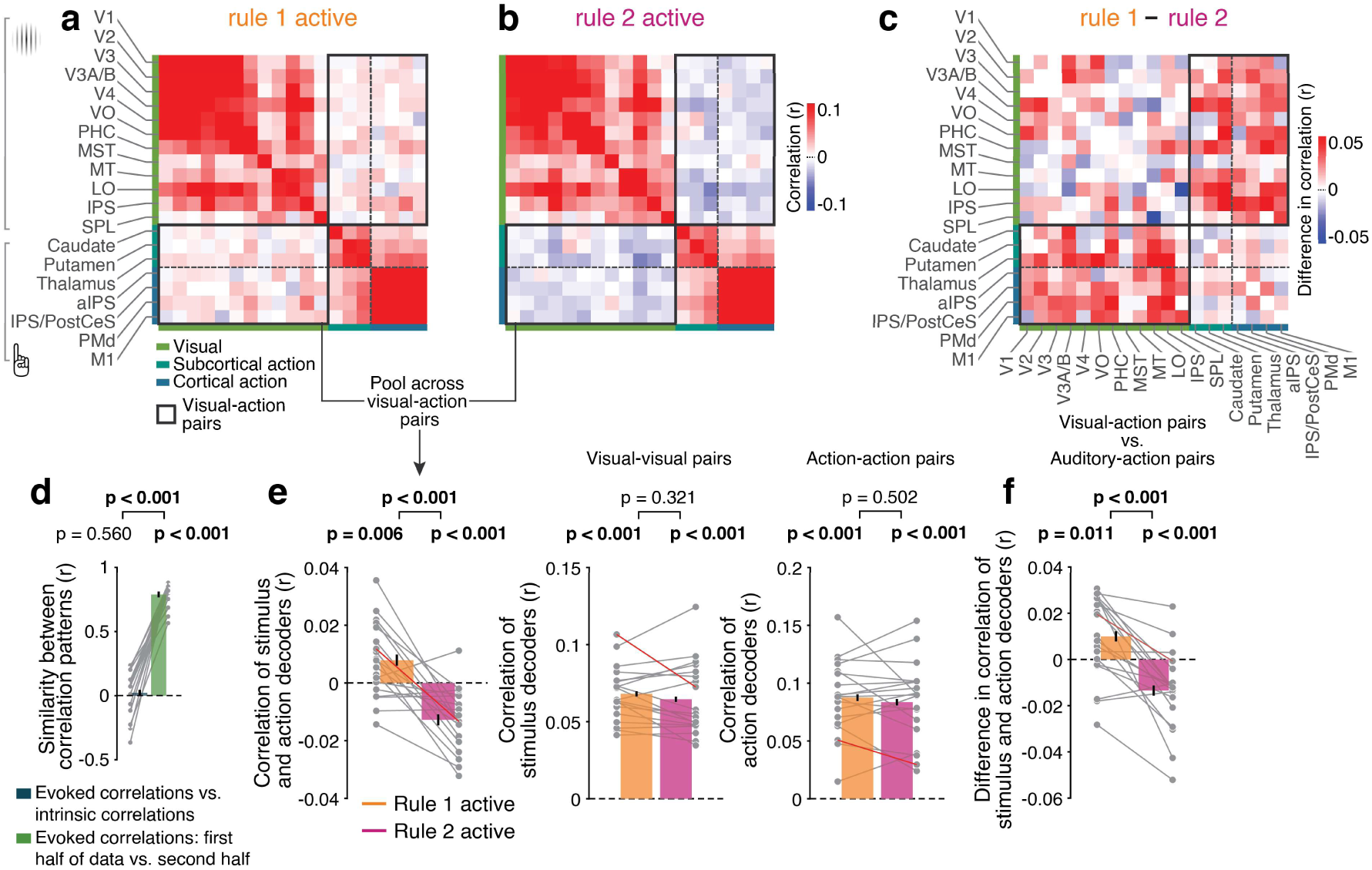
Correlations of stimulus and action codes across the sensory-motor network reflect instructed rule. **a, b.** Patterns of correlation between all pairs of decoders for stimulus orientation (visual cortical areas) and for action (subcortical and action-related cortical areas). The gray rectangle contains all pairs of stimulus and action decoders. **a.** Correlations during rule 1. **b.** Correlations during rule 2. **c.** Difference matrix between rule 1 and rule 2. **d.** Similarity of correlation patterns (across region pairs) of decoder outputs for trial-evoked responses (evoked correlations) and for the intrinsic fluctuations (intrinsic correlations, as used in main analyses). Gray lines, individual participants; bars, group average; error bars, within-participant SEM. **e.** Left: Correlations from panels a and b, collapsed across all pairs in gray rectangle. Right: Correlations from panels a and b, collapsed across all visual-visual pairs and action-action pairs (excluding values on diagonals). Gray lines, individual participants; red line, participant from Figure 2g; bars, group average; error bars, within-participant SEM. **f.** Correlations from panel e (left), relative to the correlations of the graded outputs of a stimulus orientation decoder that was trained on auditory cortex (Figure S5), and action decoder. N = 19 for all panels. All *p*-values based on permutation tests; boldface, *p* < 0.05 (FDR corrected; Table S1).

The intrinsic correlations (i.e., computed from the outputs of decoders applied to residual activity after removing evoked activity) exhibited a spatial pattern across region pairs (gray rectangle in Figure 3c) that was distinct from the correlation pattern of the feature-specific evoked responses (termed ‘evoked correlations’ in the following; Figure 3d and Figure S4). These observations, combined with our effective removal of evoked responses (Figure S3), indicate that the measured correlations were not driven by evoked responses.

A sign flip of correlations was not evident for pairs of visual cortical areas or for pairs of action-encoding areas (Figure 3c, cells outside of gray rectangle; Figure 3e, middle, right). Furthermore, when substituting the visual cortical regions for a set of control regions centered on primary auditory cortex, no significant (and substantially weaker) correlation between stimulus and action decoder outputs was present (Figure 3f; Figure S5a-g). These control analyses highlight the specificity of the result presented in Figure 3e for the co-fluctuations of population codes for visual stimulus and motor action. In sum, the active rule was encoded in patterns of spontaneous co-fluctuations of stimulus and action codes across a task-relevant sensory-motor network of brain areas.

### Inferring volatile sensory-motor mapping rules under uncertainty

Because in the task (Figure 2a) the active rule was explicitly instructed without ambiguity, we assumed that participants’ internal belief about the appropriate sensory-motor association had close-to-maximal certainty. In many real-world situations, however, this belief is subject to uncertainty; it needs to be learned from noisy and incomplete information (Miller and Cohen, 2001; Durstewitz et al., 2010). We thus developed a new variant of the task to probe the evolving formation of an internal belief about the appropriate sensory-motor mapping under uncertainty. In this task, the sensory-motor mapping rule was volatile (i.e., could undergo hidden changes) and had to be inferred from noisy sensory evidence presented in the inter-trial intervals (Figure 4a). The evidence samples were the positions of small dots presented at a high rate in a narrow range left or right from the central fixation mark. Each dot position was drawn from one of two overlapping Gaussian distributions, which corresponded to the active rule at that time. The active rule (and thus the generative distribution) could switch from one sample to the next with a low probability (hazard rate; 1/70). The trials of the primary perceptual choice task were the same as before (orientation discrimination judgement), and randomly interspersed in the evidence stream, again with long and variable intervals (Figure 4a).

**Figure 4.**
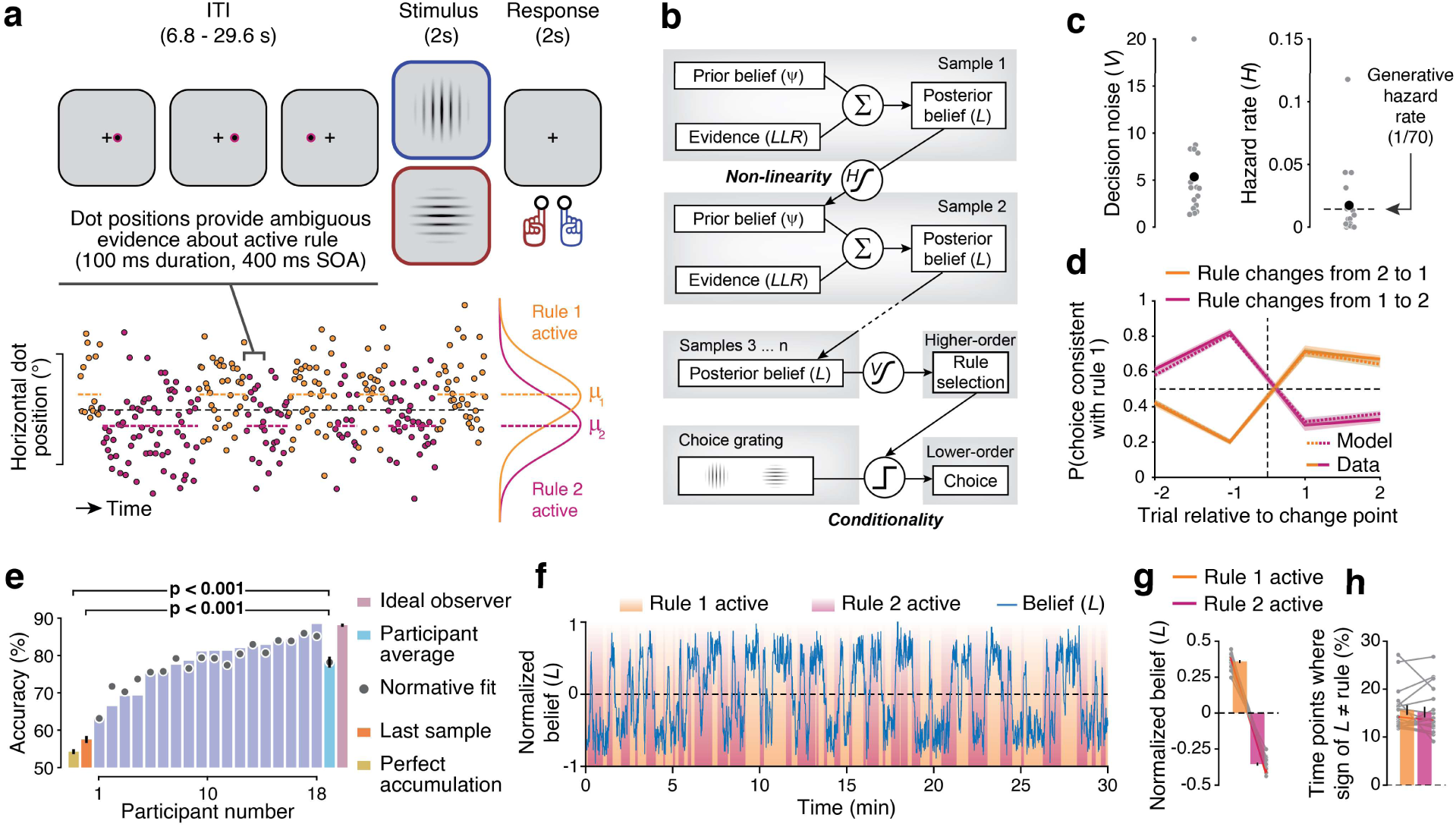
Tracking the evolution of belief about the volatile sensory-motor mapping rule. **a.** Inferred rule task. Top, example sequence of evidence samples during the ITI preceding a trial of the primary task. Evidence samples are the horizontal positions of dots presented every 400 ms. Bottom, dot positions are drawn from one of two distributions (bottom right), producing a noisy evidence stream (bottom left). The generative distribution governed the active rule and could switch unpredictably between any two samples (probability: 1/70). **b.** Schematic of normative model for rule inference and selection, coupled with the primary orientation discrimination task. See main text for details. **c.** Parameters of the best fitting model. Gray dots, individual participants; black dot, group average. **d.** Fraction of choices consistent with rule 1 surrounding changes in true (generative) rule, as produced by participants (solid lines) and best-fitting normative model (dashed lines). Shaded area, between-participant SEM. **e.** Overall accuracy of participants, best-fitting normative model, and alternative models. Error bars, SEM. **f.** Example time course of posterior belief (*L*) estimated from best-fitting model for one participant during 30 selected minutes of an experimental session. **g.** Belief averaged across data segments corresponding to true (generative) rules. Gray lines, individual participants; red line, participant from panel f; bars, group average; error bars, within-participant SEM. **h.** Percentage of time points at which the sign of belief (*L*) was inconsistent with the active rule. Gray lines, individual participants; bars, group average; red line, participant from panel f; error bars, within-participant SEM. N = 18 for all panels. All *p*-values based on permutation tests; boldface, *p* < 0.05 (FDR corrected; Table S1).

This task, called ‘inferred rule’, required participants to continuously select the currently active rule by accumulating the noisy evidence (higher-order decision) and apply the selected rule to report their decision for orientation judgment (lower-order decision). Because the generative state for the higher-order decision could undergo unpredictable and hidden state changes, perfect (i.e., lossless) accumulation of all the evidence is suboptimal, and optimal performance requires accumulating the evidence in an adaptive fashion that strikes a balance between stable evidence accumulation and sensitivity to change points (Glaze et al., 2015; Piet et al., 2018; Murphy et al., 2021).

The same participants who performed the instructed rule task also performed the inferred rule task. Again, participants reliably switched between the rules when required by the task (Figure 4d) and performed the task reasonably well overall (Figure 4e), albeit substantially below the performance level obtained in the instructed rule task (compare with Figure 2d). Note that even the ideal observer, a hypothetical agent using the normative strategy for this task (Figure 4b; see below) and with exact knowledge of the task statistics, would only perform around 88% correct on this task when given the exact same evidence sequences as presented to the participants (Figure 4e, purple).

We fitted participants’ behavior with a normative model of the adaptive evidence accumulation process. The model casts this process as dynamic belief updating (Figure 4b) (Glaze et al., 2015): Each sample of sensory evidence (expressed as log likelihood ratio, *LLR* of support for one vs. the other rule) was combined with a prior belief (also expressed as log-odds, Ψ) to form a posterior belief (*L*) for one over the other rule. The posterior was passed on to the next updating step to become the new prior that was then combined with the next evidence sample, and so on. Critically, the transformation of posterior into subsequent prior was non-linear, with a shape controlled by the agent’s estimate of the environmental hazard rate (subjective hazard rate *H*; Figure 4c). This feature rendered the model adaptive, sensitive to different levels of environmental volatility, in line with observations of human behavior across different experimental settings (Glaze et al., 2015; Murphy et al., 2021; Weiss et al., 2021). In our version of the model, this adaptive inference process for the higher-order decision (rule selection) was coupled with a lower-order decision about the grating orientation that governed the behavioral choice (Figure 4b). For simplicity, our model assumed that noise corrupted the higher-order rule selection (*V*), but not the lower-order orientation judgment (Figure 4b). We fitted the model to the participants’ choice behavior with *H* and *V* as the only free parameters.

The model fit the data well for most participants (Figure 4d-e), and it matched the data better than alternative, heuristic strategies for all participants (Figure 4e). A model variant that perfectly accumulated all samples across the run without information loss, as well as a model that selected the rule using only the last-seen sample, both resulted in significantly poorer performance than both the fitted normative model and the participants’ behavior (Figure 4d; *p* < 0.001 in all comparisons; Table S1). Furthermore, similar to findings from previous work (McGuire et al., 2014), the computational variables derived from the model, specifically the magnitude (absolute value) of the posterior belief, the magnitude of sensory evidence, and change point probability (the likelihood that a change in rule has just occurred given the sensory evidence and belief) (Murphy et al., 2021), all co-varied with fMRI activity across widespread cortical regions (Figure S6g-i).

As expected, the time-varying posterior belief (*L*) of the fitted model tracked the active rule in all participants (Figure 4f-g). Yet, the active rule and belief also deviated on a substantial fraction of time points (Figure 4h). Participants never had direct access to the active rule. So, their model-inferred belief rather than the truly active rule was the quantity that should drive their behavior in principle, and likely also did govern behavior in practice (Figure 4d-e). In the following, we thus used the model-derived belief time courses to interrogate the patterns of fMRI-based estimates of correlated variability across the sensory-motor network.

### Correlated variability of stimulus and action codes also reflects inferred rule

We assessed if the results from the instructed rule task (Figure 3) would replicate in, and generalize to, a different behavioral context. Here, we grouped all time points within the task runs into periods where the latent variable *L*, capturing participants’ hidden belief, favored rule 1 (i.e., *L* > 0) or rule 2 (*L* < 0). We then compared the patterns of co-fluctuations of stimulus and action codes across the sensory-motor network between these data segments, just as we did for the data segments corresponding to the active rule in the instructed rule task.

Again, we found a consistent difference in the sign of correlations between stimulus and action decoders between the two inferred rules (Figure 5a-c,e). As for the instructed rule task, this pattern of intrinsic correlations was dissimilar from the one of feature-specific evoked responses (Figure 5d), and absent when substituting the visual cortical regions for a set of auditory control regions (Figure 5f; Figure S5h-k). In sum, the results from Figure 5 replicate all qualitative findings from the instructed rule task (Figure 3) in independent data and show that they generalize to the (more general) behavioral context in which the agent operates under uncertainty about the correct rule.

**Figure 5.**
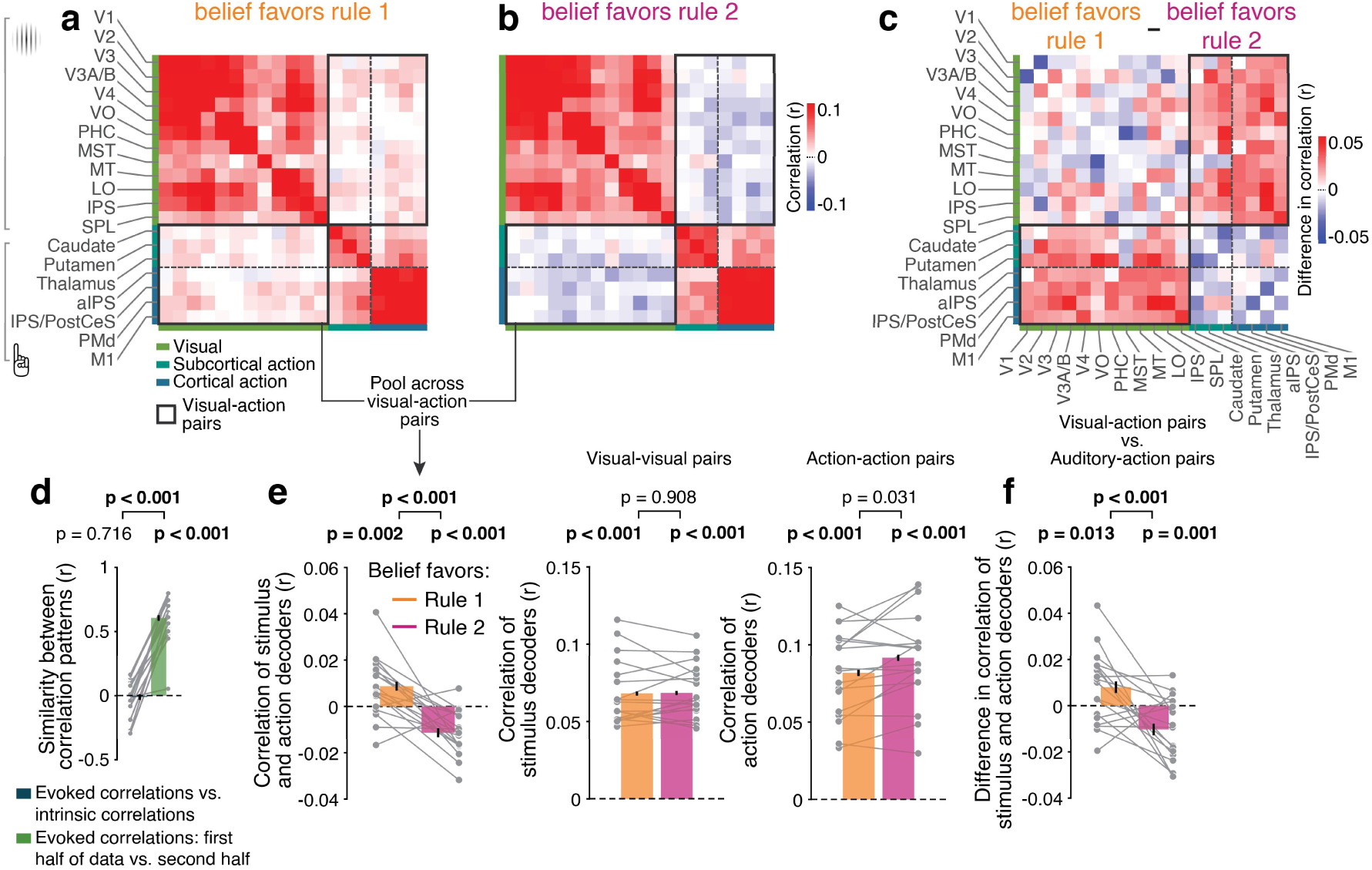
Correlations of stimulus and action codes reflect the rule inferred under uncertainty. **a-c.** Same as Figure 3a-c, but data segments for correlation based on the sign of model-derived belief (i.e., whether belief favors rule 1 or 2), rather than based on active (generative) rule. **d.** Similarity of correlation patterns of decoder outputs for trial-evoked responses (evoked correlations) and for intrinsic fluctuations (intrinsic correlations). Gray lines, individual participants; bars, group average; error bars, within-participant SEM. **e.** Left: Correlations from panels a and b, collapsed across all pairs in gray rectangle. Middle and right: Correlations from panels a and b, collapsed across all visual-visual pairs and action-action pairs (excluding values on diagonals). Gray lines, individual participants; bars, group average; error bars, within-participant SEM. **f.** Correlations from panel e (left), relative to the correlations of the graded outputs of an action decoder and a stimulus orientation decoder that was trained on auditory cortex (Figure S6). N = 18 for all panels. All *p*-values based on permutation tests; boldface, *p* < 0.05 (FDR corrected; Table S1).

### Robust decoding of instantaneous rule belief from correlated variability

Having established that rule information was reflected in the patterns of intrinsic correlations between sensory and action codes in different areas for two different behavioral contexts (instructed and inferred rule), we next asked if the rule-coding correlation patterns were similar across contexts. Indeed, the patterns of rule-coding differential correlations between region pairs in Figures 3c and 5c were similar, with pattern correlations above chance (*r* = 0.42; Figure 6a; Table S1).

**Figure 6.**
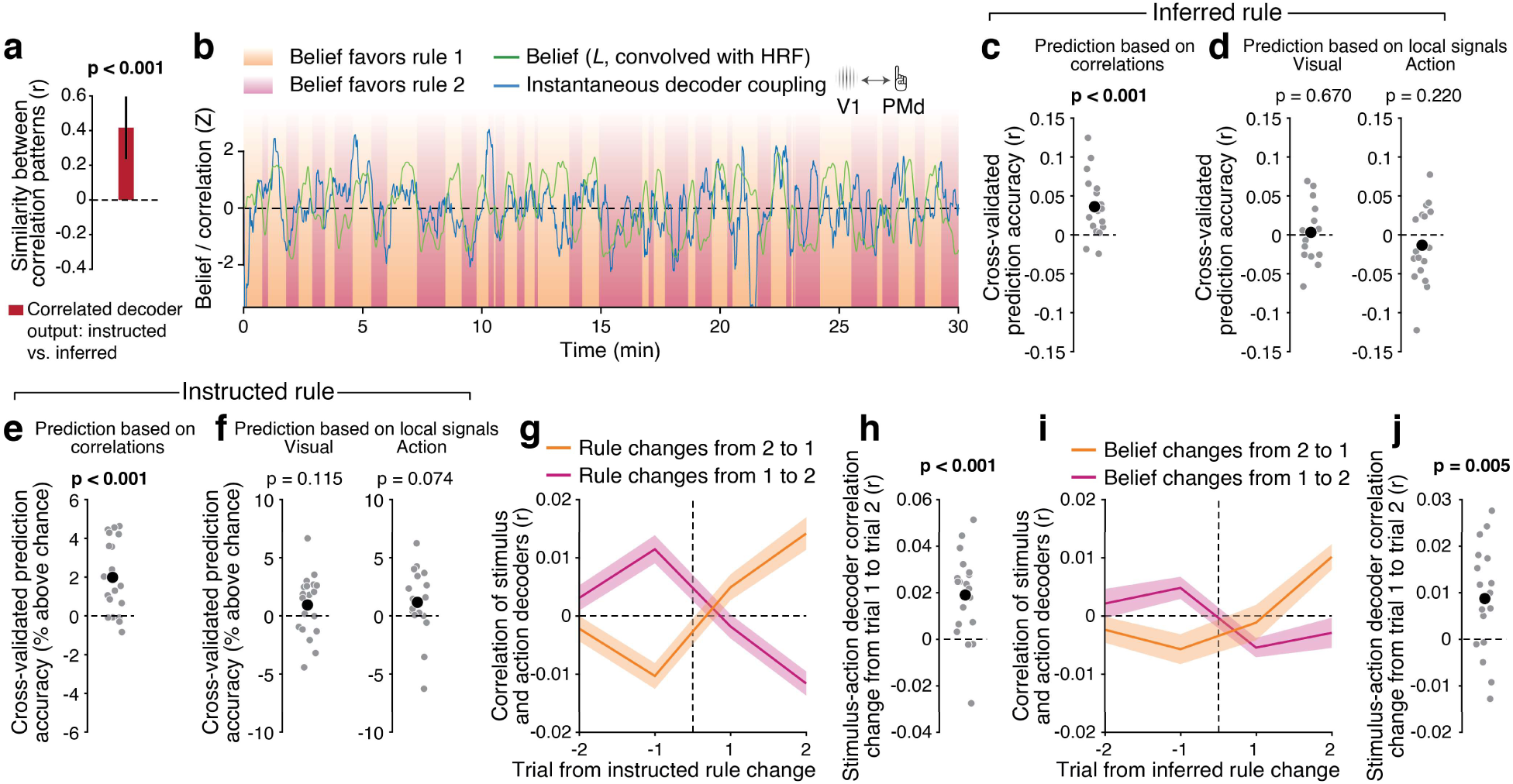
Rule belief prediction from correlations and relationship of predictive patterns across tasks. **a.** Similarity of the patterns of intrinsic decoder co-fluctuations in the instructed and inferred rule tasks (gray rectangles in Figures 3c and 5c). Error bar, 95^th^ particle of permuted null distribution. **b.** Example time courses of belief (convolved with hemodynamic response function, HRF), and time-resolved correlation between graded stimulus decoder output from V1 and graded action decoder output from PMd. The time series of time-resolved decoder correlations is smoothed for visualization purposes only. **c.** Cross-validated prediction of belief from the time-resolved coupling between stimulus and action decoders. Gray dots, individual participants; black dot, group mean. **d.** As c, but rule predictions based on either local stimulus decoder outputs or action decoder outputs, rather than their correlations. **e, f.** As c, d, but for the instructed rule task. **g.** Stimulus-action decoder correlation as function of trials from rule changes in the instructed rule task. Error bars, SEM. **h.** Change in correlation from trial 1 to trial 2 after instructed rule changes. Positive values indicate stronger correlations on trial 2 compared to trial 1. Gray dots, individual participants; black dot, group mean. **i, j.** As g,h, but for the inferred rule task. Here, the changes are defined based on sign flips of of *L*. N = 18 for panels a,c,d,i,j. N = 19 for panels e-h. All *p*-values based on permutation tests; boldface, *p* < 0.05 (FDR corrected; Table S1).

We then tested whether the correlation patterns were sufficient to predict the participants’ hidden belief about the rule at any given moment in time. The analyses reported thus far were all based on decoder correlations that were quantified across extended segments of data and categorized by the instructed or inferred rule (i.e., sign of *L*). For the inferred rule task, this approach ignored a substantial fraction of the variability of *L* that governed participants’ rule switching behavior and the associated uncertainty. This variability is illustrated in Figure 4f. In order to interrogate the relationship between rapid fluctuations of *L* and the coupling of stimulus and action codes, we estimated the latter in a time-variant fashion (Methods; see Figure 6b for an example participant). We applied a cross-validated multivariate regression model to test for robust prediction of *L* by the co-fluctuation of stimulus and action codes (Methods). Conceptually, the model contained two sets of regressors: (i) time-variant estimates of the correlations of stimulus and action decoder outputs from visual-action pairs of regions; and (ii) fluctuations of local stimulus and action decoder outputs. We trained the model (i.e. estimated regression weights) on a part of the original time series and tested its prediction accuracy on a left-out segment of the timeseries (10-fold cross-validation). This yielded significant prediction of instantaneous belief from time-resolved correlation of the stimulus- and action-decoder pairs (Figure 6c). By contrast, belief prediction was not possible based on the outputs of either the stimulus or the action decoders alone (Figure 6d).

We obtained analogous results for cross-validated rule prediction in the instructed rule task. Here, we trained a logistic multiple regression model to predict the active rule at individual time points (Methods). Mirroring the results of the inferred rule task, predictions based on the correlation of stimulus and action decoder outputs were above chance (Figure 6e), but not predictions based on the stimulus or the action decoder outputs by themselves (Figure 6f).

### Dynamics of rule-specific correlation patterns

In both tasks, the strength of the rule-specific correlations increased gradually as function of trial from the (instructed or inferred) rule change (Figure 6g-j). This contrasted with the more rapid flip of rule-consistent behavior following rule switches (compare to Figures 2c and 4d) and indicated that the switch in correlation patterns took several seconds to be instantiated. The accumulator property of the hemodynamic response could also produce a gradual increase in fMRI correlations after the switch. We verified in simulations (Figure S7) that this was not sufficient to account for the increase from trial 1 to trial 2 after the rule switch shown in Figure 6g-j. This increase (Figure 6g-j) may point to short-term plasticity involved in the mechanism underlying switches in rule-specific coupling between sensory and action-related brain regions (Miller and Cohen, 2001; Fusi et al., 2007). It is important to note that the sluggish dynamics in rule-coding correlation patterns counteracts prediction of rule belief *L* (Figure 6c-d).

### Functional significance of rule-specific correlation patterns

How did the different features of activity in the sensory-motor network assessed here relate to participants’ overt behavioral performance? Different from the instructed rule task, the inferred rule task was challenging and yielded a substantial fraction of errors (Figure 4e). Stimulus orientation decoding from patterns of evoked fMRI responses during the primary task was slightly more reliable on correct choices in certain time windows for some of the visual cortical regions (Figure S8a) and likewise for action decoding from action-related regions (Figure S8b). These subtle differences are consistent with previous findings for choice accuracy (Harvey et al., 2012) or confidence (Kiani and Shadlen, 2009; Wilming et al., 2020). Yet, critically, evoked responses enabled robust stimulus and action decoding for both correct and error trials in all regions (Figures 7a-b and S8).

**Figure 7.**
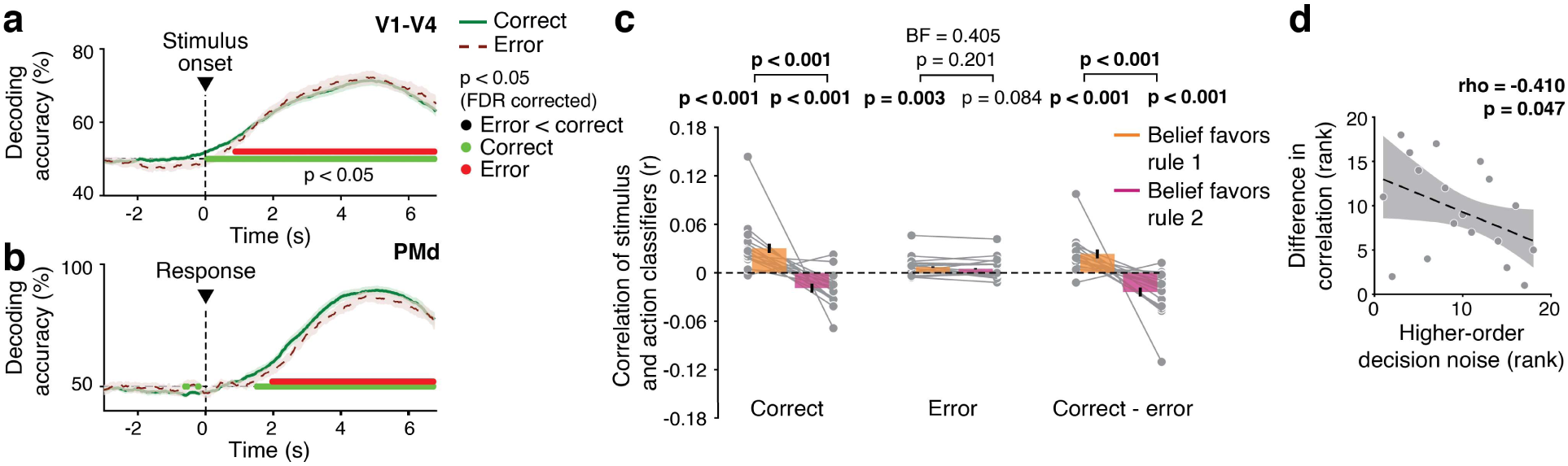
Relationship between correlated variability and behavioral performance. **a.** Accuracy of stimulus orientation decoding from evoked responses in V1-V4. **b.** Accuracy of action choice decoding from evoked responses in PMd. a-b: Shaded areas, between-participant SEM. **c.** As Figure 5e left panel, but separately for data segments surrounding correct and error trials. BF: Bayes Factor. **d.** Across-participant correlation between the model-derived noise estimates and the difference in the co-fluctuations of stimulus and action codes between the two belief states across all trials. Shaded area, 95% CI (bootstrapped) around the least squares regression line. Gray dots, individual participants. N = 18 for all panels. All *p*-values based on permutation tests; boldface, *p* < 0.05 (FDR corrected; Table S1).

The ongoing co-fluctuation between stimulus- and action-codes exhibited a relation to behavioral performance that was distinctly different from the evoked responses. While the belief-coding correlation patterns were robustly expressed on correct trials (Figure 7c, left; compare with Figure 3e), these patterns broke down on error trials (Figure 7c, middle).

Consequently, there was a marked difference between the correlation patterns on correct and error trials (Figure 6c, right). Our analysis ensured that the lower number of incorrect trials relative to correct trials could not account for the absence of the effect during errors (Methods). Furthermore, the Bayes factor provided evidence (Wetzels and Wagenmakers, 2012) for the null-effect of rule on the correlation pattern on error trials (Figure 7c).

The observation that stimulus and action decoding from evoked responses (Figure 7a-b) was above chance on error trials implies that the coupling of evoked responses flipped sign on errors compared to correct trials because a given stimulus was followed by the opposite choice in both trial types. This flip in coupling of evoked responses on errors contrasts with the absence of intrinsic rule-specific co-fluctuations on errors (Figure 7c).

Errors on the task could originate from several sources. One possible source is noise corrupting the primary orientation judgment, specifically variability of sensory responses (Arieli et al., 1996). Near-perfect performance on the instructed rule task (Figure 2d) and indistinguishable orientation decoding from early visual cortex for correct and error trials (Figure 7a) indicates that this noise source was negligible. Another source is the uncertainty about the active rule that is inherent to the task: Even a bias- and noise-free ideal observer used the wrong rule in about 12% of trials when exposed to the same evidence streams as the participants (Figure 4e, purple). A third source is suboptimality of the higher-order decision, in the form of noise or biases corrupting the evidence accumulation and/or resulting rule selection (Drugowitsch et al., 2016; Murphy et al., 2021).

Based on these considerations, we examined the relationship between individual differences in performance, and the rule-related change of correlation patterns in the sensory-motor network. While all participants performed near-perfectly in the instructed rule task (Figure 2d), they exhibited varying degrees of behavioral suboptimalities in the inferred rule task (Figure 4d; range of individual accuracies: 63% to 88%). In our model, deviations of accuracy from the ideal observer were accounted for by the individual noise parameter *V* (Figure 4b-c). Indeed, individual *V* estimates correlated with the differences in the co-fluctuations of stimulus and action codes between the two belief states (favoring rule 1 versus rule 2; Spearman’s rho = -0.631, *p* = 0.003; Figure 7d; Table S1).

Taken together, the results suggest that the flexibility of the patterns of co-fluctuation of stimulus and action codes across the sensory-motor network limited performance on our challenging task, linking it to trial-to-trial fluctuations of performance within participants (correct vs. error), as well as across-participant variation in internal noise (*V*).

### Rule and belief encoding within premotor and visual cortex

Our analyses showed that (participants’ belief about) the appropriate sensory-motor mapping rule was reflected in the correlations between local population codes for stimulus and action in the sensory-motor network (Figures 6c and 6e), but not in these local population codes themselves (Figures 6d and 6f). The latter is expected because the decoders used to estimate these population codes were each trained on the stimuli or actions only, which by design were orthogonal to the task-relevant stimulus-response association specified by the rule.

We also trained a separate set of decoders to identify putative representations of (belief about) the rule in local activity patterns in a large set of regions covering the entire cortex (Methods). Local patterns of fMRI activity afforded robust decoding of (belief about) the active rule during both the instructed rule and the inferred rule tasks in three regions: PMd (area 6d), posterior parietal cortex (area 5L, only left hemisphere), and primary visual cortex (V1, Figure 8a-c). Sub-cortical regions did not yield significant decoding of rule in either the instructed or the inferred rule task (all *p*-values > 0.075), potentially because the signal-to-noise ratio (SNR) of our fMRI measurements was lower in the sub-cortical regions compared to the cortex (temporal SNR of 163 averaged across cortical regions vs. 109 averaged across subcortical regions, *p* < 0.001).

**Figure 8.**
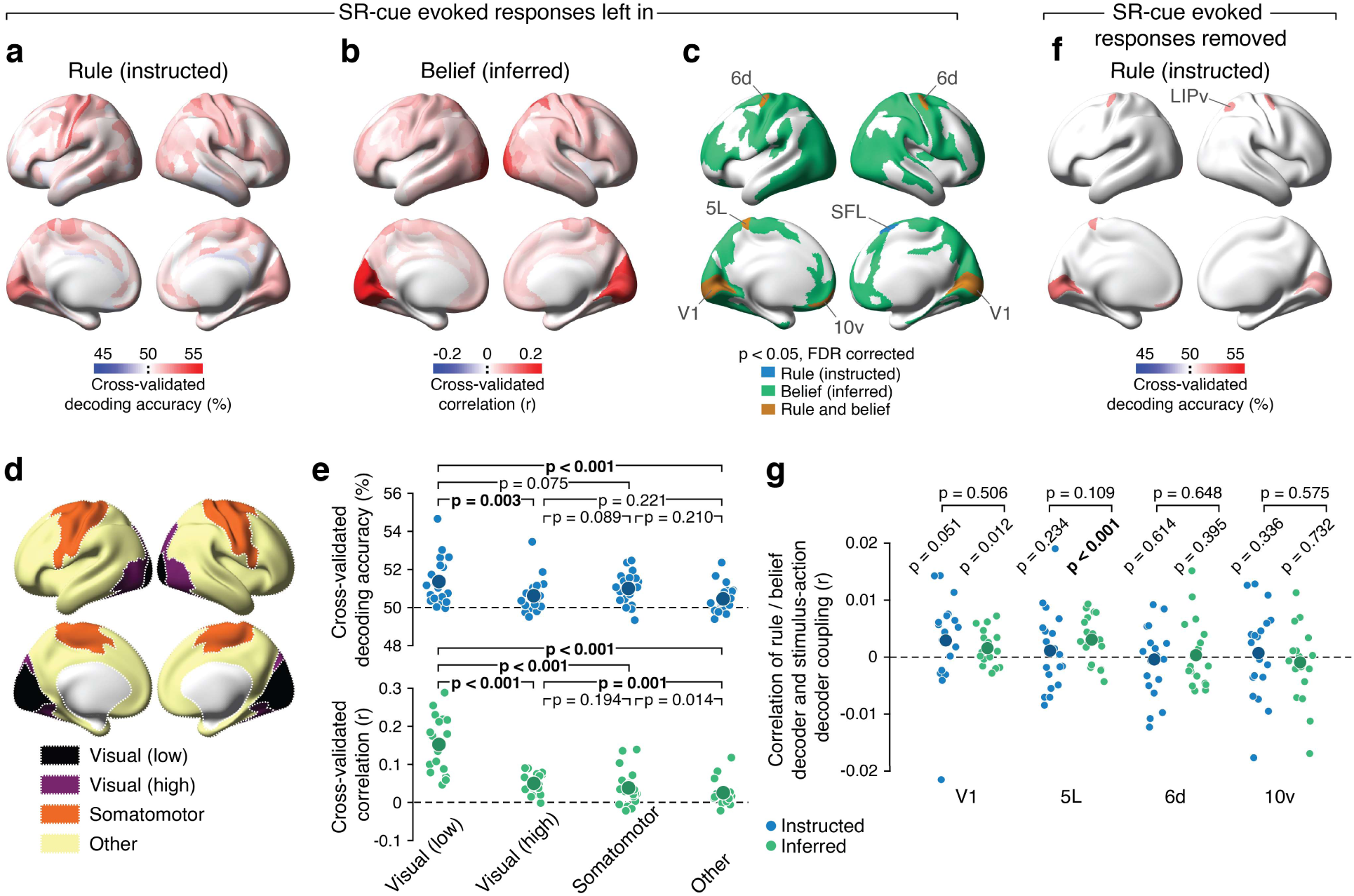
Local encoding of rule and rule belief across cortical areas. **a.** Decoding of active rule (instructed rule task) from continuous time-series data encompassing ITIs as well as orientation discrimination trials. **b.** Decoding of graded rule belief (inferred rule task) from the continuous time-series data. **c.** Overlay of areas with significant classification accuracy from a and b. **d.** Groupings of cortical areas into clusters (Methods) **e.** Comparison of decoding precision in both tasks between different sets of areas. **f.** Significant (*p* < 0.05, FDR corrected) active rule decoding (instructed rule task), after removing responses evoked by the SR-mapping cue. **g.** Correlations between rule (instructed) or belief (inferred) decoder output, and time-variant stimulus-action decoder coupling. Small dots, individual participants; Large dots, group average. ROI labels on the x-axis mark the regions on which the rule / belief decoders were trained. All *p*-values based on permutation tests; boldface, *p* < 0.05 (FDR corrected; Table S1).

Encoding of the instructed rule was more confined, restricted to the aforementioned three regions and two prefrontal cortical areas (SFL and 10v, Figure 8a,c), whereas encoding of rule belief (inferred rule) was widespread across many regions (Figure 8b-c), similar to what was observed in studies with more complex rule sets (Ito et al., 2022). The more confined effects in the instructed rule task may be due to the lower demand in tracking and implementing the active rule in this task and/or the long delays without any additional rule cues (Figure 2a).

When assessed at a coarser scale (Figure 8d, Methods), we found that lower-tier visual cortex (V1-V4) contained more rule information than other area clusters (Figure 8e). This was true for both tasks when compared with higher-tier (dorsal and ventral) visual cortex as well as a large swath of cortex containing most associative regions (’other’ in Figure 8d-e). In the inferred (but not instructed) rule task, this was also true when lower-tier visual cortex was compared with a cluster comprising premotor and somatomotor cortex (‘somatomotor’; Figure 8d-e).

Encoding of SR-mapping rules in posterior parietal and frontal cortex is consistent with primate physiology (Wallis et al., 2001; Stoet and Snyder, 2004; Sakai, 2008; Bennur and Gold, 2011), but the task-general encoding of rule information in V1 may be unexpected (but see Zhang et al., 2013; Siegel et al., 2015). One concern is that the local rule decoding effects may have been confounded by sensory responses to the visual cues that signaled the SR-mapping rule in an unambiguous (instructed rule) or ambiguous (inferred rule) fashion. In the current dataset, this concern cannot be ruled out for the inferred rule task because the time courses of cue positions and of belief state were too similar after convolution with the hemodynamic response function (mean *r* = 0.77). However, for the instructed rule task, the scenario makes two predictions, both of which were falsified. First, if responses evoked by the SR-cue drove the rule decoding (in V1 or downstream areas), then training and testing the decoder on cue-evoked responses should also yield robust rule decoding: we did not find any significant cue-encoding in the evoked responses for any cortical area. Second, removing cue-evoked responses should impair rule decoder performance. However, V1, LIPv, 5L, and 6d all exhibited clear rule encoding in the residuals (Figure 8f). Combined, these results establish that genuine rule information was present in the persistent activity of these areas, including V1.

The dynamics of the rule codes in V1 and parietal cortex (area 5L) also correlated with the dynamics of instantaneous coupling between stimulus and action codes (Figure 8e). We measured the dynamics of the local rule code in terms of the continuous output of the rule belief decoder (inferred rule) or rule decoder (instructed rule). V1 and 5L exhibited weak, but above-zero correlation for the inferred rule task (V1: mean *r* = 0.002, *p* = 0.012, 5L: mean *r* = 0.003, *p* < 0.001), where only the effect in area 5L passed FDR correction. We also found a trend for V1 in the instructed rule task (mean *r* = 0.003, *p* = 0.051). These (weak) correlations point to an interplay between local rule codes within these areas and the distributed rule codes expressed in the sensory-motor coupling patterns, consistent with rule-selective common input to the stimulus and action codes in visual and motor cortex, respectively.

### No reflection of rule in correlations of region-average fMRI signals

Our analyses of correlations between decoder outputs were based on spontaneous fluctuations of fMRI signals (Arieli et al., 1996; Fox and Raichle, 2007), over and above the evoked responses during trials of the primary choice task as well as occurring during the inter-trial intervals. A large body of neuroimaging work has used spontaneous signal fluctuations to infer intrinsic networks of brain regions and assess their dependence on neuromodulatory or behavioral state, including sensory-motor mapping rules (Fox and Raichle, 2007; Vincent et al., 2007; Honey et al., 2009; Heinzle et al., 2012; Hipp et al., 2012; van den Brink et al., 2016; van den Brink et al., 2018; Lurie et al., 2019; van den Brink et al., 2019; Zamani Esfahlani et al., 2020; Pfeffer et al., 2021). In this previous work, correlations were computed between fMRI signals from single voxels or from the voxel-average per region, thus lacking the feature-specific information entailed in the local activity patterns (Haynes and Rees, 2006; Kriegeskorte and Bandettinin, 2007).

In contrast, by correlating feature-specific signals contained in the multivariate activity patterns within each area (i.e., continuous decoder outputs) between regions (Figure 2h), our analysis approach quantified ‘informational connectivity’ (Coutanche and Thompson-Schill, 2013). A final set of control analyses tested if that aspect was indeed necessary to detect rule information in the correlation patterns. We repeated the comparison between correlations for each active rule (instructed rule) or belief about the active rule (inferred rule) as before, only this time using the voxel-average time series of the same ROIs. This did not yield any difference between rules in the inter-regional correlations (Figure 9). Thus, inter-regional correlations of average activity were uninformative about the active rule, highlighting the specificity of the rule encoding for correlations of graded decoder outputs.

**Figure 9.**
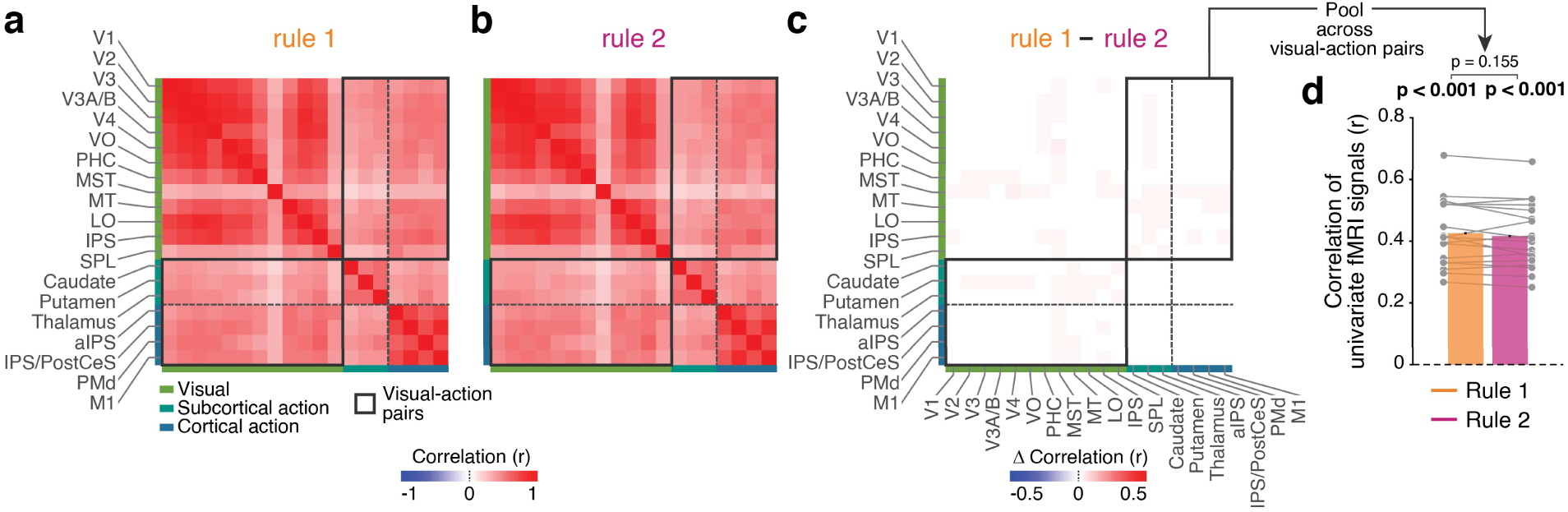
Standard functional connectivity analysis based on region-average signals. **a.** Inter-regional activity correlation when rule 1 was active / belief favored rule 1. **b.** Same as panel a, but for rule 2. **c.** The difference between rule 1 and rule 2. **d.** Inter-regional correlation strength for the edges in panel c that link the visual and motor ROIs. In contrast to the correlation of graded outputs of stimulus and action decoders, this analysis yielded no sign flip between mapping rules. N = 19 for all panels. Gray lines show the individual participants. Bars, group mean. Error bars, within-participant SEM. All *p*-values based on permutation tests; boldface, *p* < 0.05 (FDR corrected; Table S1).

## Discussion

Our findings provide new empirical constraints for a mechanistic framework of flexible, context-dependent sensory-motor transformations. Most theoretical studies of this issue have focused on the dynamics of neural population activity within individual cortical regions (Fusi et al., 2007; Mante et al., 2013; Rigotti et al., 2013; Saez et al., 2015; Mastrogiuseppe and Ostojic, 2018; Yang et al., 2019) (but see: Wang and Yang, 2018). Likewise, most experimental studies have assessed the information about such task rules that is contained in the activity of individual regions (Wallis et al., 2001; Stoet and Snyder, 2004; Haynes et al., 2007; Sakai, 2008; Bode and Haynes, 2009; Bennur and Gold, 2011; Zhang et al., 2013; Cole et al., 2016). Consequently, it has remained elusive how the large-scale network of brain regions that is involved in a given sensory-motor transformation is configured for a given task set (Shadlen and Kiani, 2013; Wang and Yang, 2018). We here developed an approach to tackle this issue. Our approach uncovered that sensory-motor mapping rules manifest in the patterns of correlated intrinsic variability of stimulus and action codes within this network.

Our approach is based on the insight that the intrinsic correlation structure of neural activity reflects the architecture of functional networks (Gerstein and Perkel, 1969; Büchel and Friston, 2000). Previous neuroimaging work has exploited this principle to infer coarse-grained functional brain networks (Fox and Raichle, 2007; Vincent et al., 2007; Honey et al., 2009; Hipp et al., 2012). Our approach combined this principle with the assessment of feature-specific population codes that are expressed in the fine-grained activity patterns within each brain region. Indeed, this combination was critical for revealing the encoding of (beliefs about) stimulus-response mapping rules in intrinsic co-fluctuations across regions: rule-information was specifically contained in the correlations between stimulus- and action-specific activity patterns in different brain regions (Figures 2-6), not in the correlations of the same regions’ mean activity (Figure 9). The distributed form of rule encoding we revealed here was proportional to behavioral accuracy (Figure 7) and co-existed (and interacted) with local rule codes expressed within individual cortical areas (Figure 8).

Correlations of ongoing activity are an important property of collective neural dynamics (Wang, 2022) but have not yet been explored in computational analyses of flexible input-output mapping. One study of perceptual decision-making, unrelated to sensory-motor mapping, has analyzed correlations within and between sensory and motor modules of a recurrent neural network that was trained on a visual evidence accumulation task (Pinto et al., 2019). Our findings may inspire future studies to train such large-scale and modular networks to solve rule switching problems. The correlation patterns between the sensory and action codes in these various modules can be analyzed with the same approach we present here. These correlation patterns could prove to be a powerful marker of the flexible switching between distinct ‘communication subspaces’ of interacting neural populations situated in different brain regions (Shenoy and Kao, 2021).

One line of work on sensory-motor mapping rules has developed the notion that neural populations in association cortex act as local switches that are transiently activated by the conjunction of stimulus and rule, and route sensory signals from sensory regions to action-related regions in a rule-dependent fashion (Cocuzza et al., 2020; Kikumoto and Mayr, 2020; Ito et al., 2022). This idea builds on the observation of mixed selectivity of neurons in prefrontal cortex with responses that combine several stimulus properties and task cues in a non-linear fashion (Rigotti et al., 2013). Yet other work has focused on the role of disinhibitory circuit motifs for the dynamic gating of specific inter-area pathways (Wang and Yang, 2018). These views are not mutually exclusive with our current results. Rather, our results complement them with a perspective on the large-scale cortical correlation structure.

A few previous studies have probed the neural bases of switching between sensory-motor mappings with inherent asymmetries (Heinzle et al., 2012; Sarafyazd and Jazayeri, 2019; Duan et al., 2021), such as pro- and anti-saccade tasks (Munoz and Everling, 2004). Here, the ‘pro-rule’ requires using a neural default pathway between matching positions within the spatial maps in sensory and action-coding brain regions, whereas the ‘anti-rule’ requires overriding this default pathway. By contrast, in our task, the mapping from the feature space of visual orientation to the space of motor action required a selective routing of signals from populations of visual cortical neurons to action-coding neural populations, without any asymmetry between the rules, as evident in the highly symmetric behavioral performance under both rules (Figure 2c-e). Recent work has begun to illuminate the local circuit mechanisms underlying the switches between asymmetric rules in prefrontal cortex (Sarafyazd and Jazayeri, 2019) and the brainstem (Duan et al., 2021). It may be instructive to compare large-scale patterns of rule-selective connectivity in cortical sensory-motor pathways between asymmetric and symmetric rule settings.

Our findings extend recent work showing that humans (Glaze et al., 2015; Filipowicz et al., 2020; Murphy et al., 2021; Weiss et al., 2021) and rats (Piet et al., 2018) can accumulate perceptual evidence in volatile environments in an approximately normative fashion. In all these studies, the evidence accumulation process pertained to perceptual judgments about the state of the sensory environment. By contrast, in our task, the evidence accumulation informed a higher-order decision: the selection of an internal task model (i.e., the appropriate mapping rule) used to report the outcome of a lower-order perceptual decision. The observation that the same normative process can explain human behavior also in the current context generalizes these previous findings to higher-order decision-making and establishes the adaptive nature of the human inference machinery.

Two previous studies have shown that humans and non-human primates can effectively implement hierarchical decision processes in volatile environments (Purcell and Kiani, 2016; Sarafyazd and Jazayeri, 2019). In these studies, participants needed to infer possible changes in the sensory-motor mapping rule from negative choice outcomes after a challenging perceptual judgment that was subject to uncertainty. Their behavior could be explained by models that integrated previous choice outcomes with the expected accuracy of perceptual judgments to infer rule switches. By contrast, in our inferred rule task, trial-by-trial outcome feedback was not provided, and uncertainty was restricted to the rule inference problem. Thus, our findings extend the notion of hierarchical decisions (in our case: about rule and stimulus orientation) to a setting that requires a different inference strategy, in which participants integrated noisy cues on a sensory dimension that was orthogonal to the one for the primary task. The coupling of integration processes at different hierarchical levels seems to be a key principle used by the brain to generate adaptive behavior in uncertain environments.

The main insight provided by the current study bears striking resemblance to recent analyses of task-dependent correlated variability between individual neurons in animals. We found that fine-grained patterns of inter-area correlations between local feature-selective activity patterns reflect beliefs about sensory-motor mapping rules and predict behavioral performance. Cellular-resolution recordings have established task-dependent patterns of correlated variability in visual cortex (Cohen and Newsome, 2008; Haefner et al., 2016; Bondy et al., 2018) and stronger single-neuron noise correlations on correct compared to erroneous choices in parietal cortex (Valente et al., 2021). These studies have commonly assessed activity patterns within individual cortical regions and did not study the dynamics of task-specific correlated variability as we did here. Furthermore, the relationship between single-neuron noise correlations and intrinsic fMRI correlations is poorly understood and may be complex (Zhang et al., 2020). Even so, the analogy of findings points to a common functional principle that may underlie adaptive patterns of correlated neural variability at different scales – for example, top-down signaling from rule-encoding regions (Miller and Cohen, 2001; Sakai, 2008), neural sampling (Fiser et al., 2010; Haefner et al., 2016), or large-scale attractor dynamics (Wang, 2022). It is tempting to speculate that correlated neural variability may be a general format the brain employs for encoding contextual variables: Different from sensory, motor, or even cognitive variables (e.g., value or symbolic meaning), such contextual variables do not need to be ‘read out’ from any downstream neural population, but rather control the information flow through the sensory-motor network.

## Acknowledgements

We thank Joshua Gold, Matthew Nassar, Sander Nieuwenhuis, Srdjan Ostojic, and Stefano Panzeri for helpful discussion, and Jürgen Finsterbusch for help with preparation of MRI sequences. This work was funded by the Deutsche Forschungsgemeinschaft (DFG, German Research Foundation) – DO 1240/3-1 (THD), DO 1240/4-1 (THD), and SFB 936 - 178316478 - A6 (CB), A7 (THD) & Z3 (THD).

## Author contributions

**RLvdB:** Conceptualization, Methodology, Software, Formal analysis, Resources, Data Curation, Writing – Original Draft, Writing – Review & Editing, Visualization. **KH:** Methodology, Software, Formal analysis, Data Curation, Writing – Review & Editing. **NW:** Conceptualization, Methodology, Software, Formal analysis, Investigation, Data Curation, Writing – Review & Editing, Supervision. **PRM:** Methodology, Software, Formal analysis, Writing – Review & Editing. **CB:** Conceptualization, Writing – Review & Editing. **THD:** Conceptualization, Methodology, Resources, Writing – Original Draft, Writing – Review & Editing, Visualization, Supervision, Project administration, Funding acquisition.

## Materials and Methods

### Participants

A total of 22 participants (median age 27, range 21 – 44, 8 male) took part in our experiment. All participants gave written informed consent and the study was approved by the ethics committee of the Hamburg Medical Association. All participants were healthy individuals with normal or corrected vision recruited via the recruitment pool of the Department of Neurophysiology and Pathophysiology of the University Medical Center Hamburg-Eppendorf. Exclusion criteria included a current or past diagnosis of mental or neurological illness, use of illegal substances, above-average consumption of alcohol as well as non-compatibility with the MRI-scanner.

The experiment comprised three sessions, one behavioral training session and two sessions in the MRI-scanner. All but one participant completed all three sessions. This one participant, plus another two were excluded from MRI analyses, the latter two because of a failure to record pulse and respiration. One further participant was excluded from the inferred rule task (see below) because of an error in logging the response data. Thus, 19 (instructed rule) and 18 (inferred rule) out of 22 tested participants were included in the analyses presented here.

Participants were remunerated with 10 Euros per hour, 10 Euros for completing all three sessions, and a variable bonus, the amount of which depended on task performance across all three sessions. The maximum bonus was 30 Euros.

### Behavioral tasks

Participants performed two different versions of a hierarchical decision-making choice task, which combined the selection of a changing sensory-motor (SR) mapping rule (higher-order decision) with a basic visual orientation discrimination judgment (lower-order decision). The SR-mapping rule defined the correct mapping from visual orientations to motor responses (button press with either left or right index finger; Figure 1a). Participants needed to apply the selected rule to report their orientation judgment in order to obtain a monetary reward (5 cents per correct choice in the ‘inferred rule’ and 3.5 cents in the ‘instructed rule’ task version). The lower-order decision was the same in both versions of the task. Stimuli for this decision were large contrast gratings of either vertical or horizontal orientation (see *Stimuli and Procedure* for details).

The two different versions of the task, called ‘instructed rule’ or ‘inferred rule’, differed only in the difficulty of the higher-order decision (i.e., rule selection). During ‘instructed rule’ runs, visual SR-mapping cues, presented transiently (duration: 400 ms) during the inter-trial-intervals (ITIs) every two trials, instructed participants unambiguously that one of the two possible stimulus-response mapping rules would be active on the next pair of trials. The active rule alternated every pair of trials. Thus, in instructed rule runs, the selection of the SR-mapping rule was both unambiguous and predictable. ITIs for the lower-order decision were long and variable (uniform: 4-20 s).

In ‘inferred-rule’ runs, by contrast, the active rule had to be continuously inferred from a sequence of noisy sensory evidence samples and it could undergo hidden changes at any moment during each run. Participants monitored a rapid stream of evidence samples: small dots that were flashed briefly (100 ms, 400 ms SOA) around the horizontal meridian. The sample positions were drawn from one of two generative distributions: two Gaussians with equal standard deviation (σ left = σ right) and different means symmetric around fixation (|µ left| = |µ right|). The generative distribution at any moment governed the active rule, and it could change from one sample to the next with a low probability (hazard rate) of 0.0143. At variable ITIs (uniform: 6.8 – 29.6 s), the stimulus for the lower-order decision appeared, prompting participants to report their orientation judgment. The correctness of the response depended on both the selection of the correct rule and the correct orientation judgment. Thus, in inferred rule runs, participants needed to continuously infer and select the rule that was most likely to be active at any time. To this end, they needed to integrate the noisy rule evidence over time. Finally, they needed to apply the selected rule on the next trial of the lower-order decision (orientation discrimination).

Participants were instructed at the beginning of each block, which distribution corresponded to which rule. The relationship between the two generative distributions and the response rules stayed constant across experimental sessions.

### Stimuli and procedure

All stimuli were created using Matlab and the Psychophysics Toolbox Version 3 (Brainard, 1997) and presented on a medium grey background. The fixation mark, presented throughout the entire run in the center of the screen, was a white symmetric cross with a length of 0.51° of visual angle and a thickness of 0.05°. The grating stimuli for the lower-order decision were circular, achromatic Gabor patches with full contrast and truncated at an inner eccentricity of 2.5° and an outer eccentricity of 13.85°. The spatial frequency was fixed at 1.2 cycles/°, whereas orientation (the discriminandum) varied randomly from trial to trial and was either vertical or horizontal.

In the MRI-scanner, stimuli were presented on an MRI-compatible LCD-screen with a resolution of 1920×1080 pixels at a refresh rate of 60 Hz. The screen was positioned at an approximate distance of 60 cm and viewed through a surface mirror that was mounted on top of the head coil. In the training session in the psychophysics lab stimuli were presented on a VIEWPixx monitor (VPixx Technologies, Saint-Bruno, Quebec, Canada) with the same resolution and refresh rate as the monitor in the MRI-scanner. The grating stimuli spanned the full height of the projection screen. Participants reported their decisions by button presses with their left or right hand. To this end, they used two MRI-compatible button interfaces (Current Designs, Philadelphia, Pennsylvania, USA) in the scanner and a keyboard in the psychophysics lab. At the end of each run, participants received feedback regarding their overall performance in that run, in the form of (i) the percentage of correct choices, (ii) the monetary reward gained in the run, and (iii) their total monetary reward accumulated across the whole experiment up to that moment.

The two MRI sessions of each participant took place on the same testing day with a break of 105 minutes in between sessions. The training session took place 1-2 days before the MRI sessions. During the training session, participants first performed a run in which the cue that instructed the rule was continuously presented. Next, participants performed one instructed rule run, in which the participants were still informed about the correct rule before each trial, but the cue was only transiently presented, and had to be remembered upon trial presentation.

Afterward, participants performed five runs of the final task version that was used during the main experiment (i.e., MRI sessions). Before the onset of each run, participants were shown a visualization of the mapping of generative distributions to rules, in order to avoid error trials resulting from false rule association.

In each MRI session, we first ran three blocks of retinotopic mapping (not used for the current article). Then, participants performed three blocks of the inferred rule task, which on average lasted 609 s (SD = 0.54 s) and included 36.0 choices (SD = 1.94). Thereafter, participants performed two instructed rule runs, which lasted on average 604 s (SD = 6.40 s) and included on average 56.5 choices (SD = 1.44). As during the training session, participants were reminded of the correspondence between generative distributions and active rule.

### MRI data collection

All MRI data were collected with a Siemens Prisma 3T MRI scanner with a 32-channel head coil. During task performance on each of the two MRI sessions, we collected 5 runs (3 inference, 2 instructed) of T2*-weighted EPI data (Flip angle: 70; TR: 1.9 s; TE: 28 ms; FOV: 112 x 112 x 62 slices of 2 mm isotropic voxels; 328 volumes). Cardiac pulsation and breathing during task performance were recorded using a pulse oximeter and pneumatic belt. At the end of the second session, we collected a T1 high resolution anatomical scan (MPRAGE; Flip angle: 9; TR: 2.3 s; TE: 2.98 ms; FOV: 192 x 240 x 256 slices; 1 x 1 x 1 mm) for registration purposes. On both MRI sessions, we also collected B0 field homogeneity scans for field distortion correction purposes (Phase image: flip angle: 40; TR: 0.678 s; TE: 7.88 ms; Magnitude image: flip angle: 40; TR: 0.678 s; TE: 5.42 ms).

### Eye tracking and analysis of eye position

Pupil diameter and gaze position were recorded at 1000 Hz with an MRI-compatible EyeLink 1000 eye tracker and calibrated with a 9-point fixation routine. Blinks and missing data segments were linearly interpolated in the eye position data. To create 2-dimensional histograms of eye position, eye position data were sub-divided into 192 equally spaced bins along the X-dimension and 108 bins along the Y-dimension. These bins were proportional to the dimensions of the stimulus presentation screen and covered its extent in degrees visual angle. Histograms were made separately per active rule (instructed rule task) or sign of belief (inferred rule task) and submitted to bin-wise comparisons between rules using permutation testing.

### Behavioral modeling

#### Normative model

The normative solution for the higher-order decision problem (rule selection) in the inferred rule task was cast as a dynamic belief updating process (Glaze et al., 2015). In this model, belief *L* was expressed in units of log-odds, for one possible state (rule 1) versus the alternative possible state (rule 2). For each evidence sample *X*_n_, the posterior belief *L*_n_ was computed by combining the log-likelihood ratio *LLR*_n_ associated with that sample, with the prior belief ψ_n_. The key feature rendering the model adaptive in volatile environments (i.e., possibility of hidden state changes) was a non-linear transformation of the posterior from each updating step into the prior for the next step, dependent on the subjective hazard rate *H*:

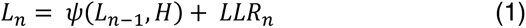

Here, *LLR*_n_ was the logarithm of the ratio of the likelihoods of the sample *X*_n_ to be observed under each of the two states, whereby the states corresponded to the generative distributions. *LLR*_n_ could also be expressed as the difference of the logs of the likelihoods:

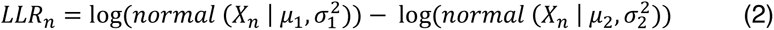

The non-linear transformation of posterior into the next prior, ψ(*L*_n-1_, *H*) was given by:

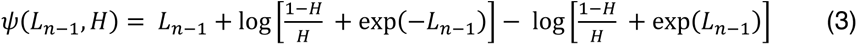

This transformation discounted the previous posterior as a function of the participant’s estimate of the environmental hazard rate *H* (i.e., probability of a change in generative distribution) and constituted the key difference to previous evidence accumulation schemes (Bogacz et al., 2006). When *H*=0, ψ= *L*_n-1_, resulting in perfect accumulation (no discounting). When H=0.5, the three righthand terms in equation (3) cancel out, so that ψ = 0 and *L*_n_ = *LLR*_n_ (complete discounting). Thus, *H* balanced the impact of new evidence and prior belief on the current belief and thereby controlled the tradeoff between stable evidence accumulation and sensitivity to change-points. The prior was initialized at the start of each run (ψ_1_ = 0) and then evolved throughout the run according to eq. 3.

We assumed that rule selection for each sample *n* was based on *L*_n_ corrupted by a Gaussian noise term *V*, such that the probability 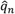 of selecting rule 1 was computed as:

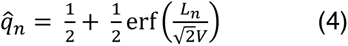

This rule selection probability then determined the probability of each action choice (left or right) upon presentation of a vertical or horizontal grating on trial *trl*. Specifically, the probability of choosing a left response 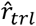 was computed as:

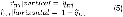

where 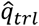 corresponds to the estimated rule selection probability after accumulating the sample directly preceding the grating presentation on trial *trl*.

#### Model fitting

We assumed that the lower-order decision (orientation judgment) was noise-free so that behavioral performance was only limited by the higher-order decision (rule selection) and a correct observed behavioral choice on a given trial implied the latent selection of the correct rule. Furthermore, we assumed no biases other than a potentially biased internal representation of the true hazard rate. Consequently, *H* and *V* were the only free parameters in our model fits to participants’ data.

Following previous work (Glaze et al., 2015; Murphy et al., 2021), we fitted the model separately to each participant’s data, by minimizing the cross-entropy between the choices of that participant and the model:

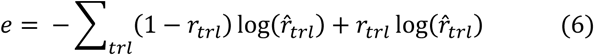

where *r_trl_* was the participant’s choice on trial *trl* (left = 1, right = 0) and 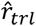 was the model choice probability on the sample *n* that corresponded to the onset of the same trial (eq. 5). The sum of *e* was minimized via particle swarm optimization. We set wide bounds on all parameters and ran 300 pseudorandomly initialized particles for 1,500 search iterations.

Having fit the model, we computed the participant- and session-specific time courses of *L*_n_ and *LLR*_n_ for analysis of the fMRI data (see *MRI data analysis* below).

#### Alternative models

We also computed the performance of three alternative models without noise: i) selecting the rule based on perfect evidence accumulation across the run (i.e., *H* = 0), ii) selecting the rule based on only the last evidence sample (i.e. *H* = 0.5), and iii) the ‘ideal observer’, which used the normative belief updating process described by eqs. 1-3 with the true generative *H* (i.e.; *H* = 1/170). We computed the performance of these models for the same evidence streams presented to the participants.

#### Model-derived change-point probability

We used the model fits to compute change-point probability (CPP) associated with each evidence sample, a computational quantity that both the normative belief updating process and human participants are sensitive to in volatile decision-making contexts such as ours (Murphy et al., 2021). CPP was the posterior probability of a change having occurred, given *H*, the previous posterior and the new evidence sample. CPP was computed as follows (Murphy et al., 2021):

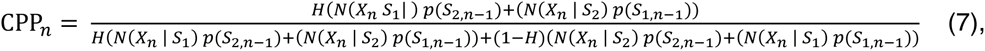

where *S*_1_ and *S*_2_ denoted the two generative distributions with mean and variance μ_1_, 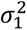 and μ_2_, 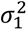.

CPP_n_ was used for univariate regressions against fMRI data (see *MRI data analysis* below).

### MRI data analysis

#### Preprocessing and physiological noise correction

We used tools from the FMRIB Software Library for preprocessing of the MRI data (Smith et al., 2004; Jenkinson et al., 2012). EPI scans were first realigned using MCFLIRT motion correction and skull-stripped using BET brain extraction. We used B0 unwarping to control for potential differences in head position each time the participant entered the scanner and resulting differences in geometric distortions in the magnetic field. The B0 scans were first reconstructed into an unwrapped phase angle and magnitude image. The phase image was then converted to units rad/s and median-filtered, and the magnitude image was skull-stripped. We then used FEAT to unwarp the EPI images in the y-direction with a 10% signal loss threshold and effective echo spacing of 0.279998. Following field-map correction, the EPI data were high-pass filtered at 100s, pre-whitened, and corrected for physiological noise using retrospective image correction (RETROICOR) (Glover et al., 2000).

RETROICOR was applied by assigning phases of the cardiac and respiratory cycles to each volume in the EPI time series, and removing them from the data. To this end, pulse oximetry (i.e., cardiac) and respiratory time series were first down-sampled from 500 Hz to 100 Hz. Next, the pulse oximetry data were bandpass filtered between 0.6 and 2 Hz, and the respiration data were low-pass filtered at 1 Hz, using a two-way FIR filter. We extracted peaks in each time series corresponding to maximum blood oxygenation and maximum diaphragm expansion, which were used to construct 34 slice-specific time-series (4^th^ order harmonics for cardiac cycle; 4^th^ order harmonics for respiration cycle; 2^nd^ order harmonics for cardiac-respiration interactions; 2^nd^ order harmonics for respiration-cardiac interactions; 1 heart rate time series; 1 respiratory volume time series). These time-series were used to estimate cardiac and respiratory effects from the EPI time series using multiple linear regression. All further analyses described below proceeded on the residuals from this regression.

Subsequently, slice time correction was applied and the EPI data were co-registered with the anatomical T1 image to 2 mm isotropic MNI space. To this end, we used FLIRT and subsequently FNIRT for maximal anatomical alignment. No spatial smoothing was applied in order to preserve high-spatial frequency information in the functional data. All critical analyses presented in this paper were applied at the level of regions of interests (ROIs, see next section).

#### Delineation of ROIs

We analyzed the fMRI data for a large set of ROIs, defined based on functional and anatomical properties based on two published MRI-based atlases (Wang et al., 2015; Glasser et al., 2016), our lab’s own previous work (de Gee et al., 2017), and the Harvard-Oxford structural atlas (https://neurovault.org/collections/262/), the latter only for sub-cortical ROIs. The primary set of ROIs that we used for all analyses presented in the main paper, selected to span the sensory-motor pathway underlying the lower-order decision, is defined in Table 1. In addition, we used the complete multi-modal parcellation from the human connectome project (HCP-MMP1.0) (Glasser et al., 2016) for supplementary analyses (see below: *Decoding of rule or belief from local activity patterns*).

**Table 1.**
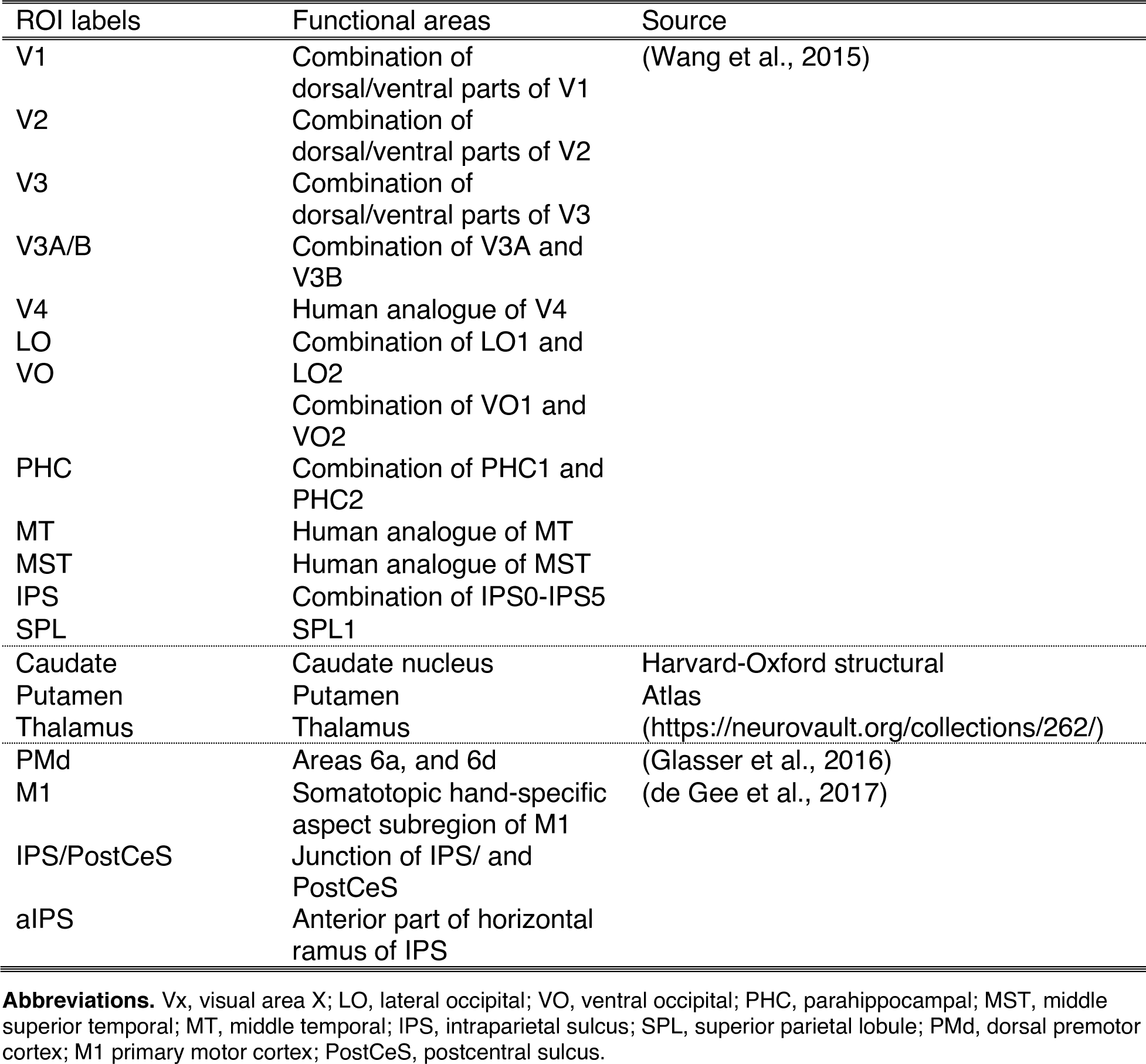
ROI definition.

##### Visual cortical areas

Our primary ROI set used an established parcellation of retinotopically organized visual field maps (Wang et al., 2015) to delineate visual cortical areas, some of which were further combined into clusters. For all analyses referring to ‘early visual cortex’ (Figs. 2 and 6), we further combined areas V1-V4 into a single ROI.

##### Action-related areas

We used our previous data (de Gee et al., 2017) to define the hand-specific somatotopic subregion of M1 as well as two posterior parietal regions (non-overlapping with the parietal visual field maps), all of which exhibited robust lateralization of activity related to the planning and execution of right versus left hand button presses across a range of previous fMRI and magnetoencephalography (MEG) studies (de Gee et al., 2017; Wilming et al., 2020; Murphy et al., 2021).To this set of action-related ROIs, we added dorsal premotor cortex from the HCP-MMP1.0 parcellation, as well as the caudate nucleus, putamen, and thalamus from the Harvard-Oxford structural atlas, based on the observation that those exhibited robust hand movement-selective activity.

#### Deconvolution of evoked fMRI responses

We used deconvolution to estimate evoked responses time-locked to specific experimental events (visual grating stimuli, button presses, or SR-mapping cues) without making any assumptions about the shape of the hemodynamic response (Dale, 1999). These trigger events were not synchronized with the EPI acquisition. Thus, we first up-sampled the fMRI data by creating a 1 x *N* vector *S* at 50 times the sampling rate, where *N* denoted the number of up-sampled time points. *S* was filled with value 1 at the time point of the trigger event, and 0 in all other positions. *S* was then iteratively staggered forward by one sample *P* times to yield an *N* x *P* design matrix *X*, with value 1 along the diagonals locked to stimulus onset. *P* was set to 396, equivalent to ∼15 seconds. The estimated response *R* (a total of *P* samples) was then obtained via:

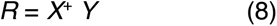

where ^+^ denotes pseudoinverse. *Y* was an *N* x 1 up-sampled, linearly detrended and z-scored time series (single voxel or ROI-average, see below).

#### Estimation of individual hemodynamic responses functions (HRFs)

Individual HRFs were estimated non-parametrically using deconvolution (see previous section) of the evoked V1 responses to the grating stimulus presentation. For this analysis, time series were first averaged across all voxels within V1. The resulting response was baseline-corrected by subtraction of the first sample from each sample, and it was normalized to unit height. This response estimate was used as an empirically derived HRF model that was convolved with the regressors in all general linear model (GLM) analyses reported below.

#### Decoding of stimulus and action from patterns of evoked responses

Decoding of stimuli (horizontal versus vertical orientation) and action (left versus right hand button press) was performed on the data of individual participants using a support vector machine (SVM) classifier with linear kernel (cf. Wilming et al., 2020). For each ROI, we selected the 300 voxels that exhibited maximum fMRI-responses to the grating stimuli (selection for stimulus decoding) or maximum lateralization of fMRI-responses during button presses (for action decoding). Maximal stimulus- and action-evoked fMRI responses (response lateralization for action) were determined based on a GLM of stimulus and action onto the voxel-level fMRI data (see *Whole brain GLMs* below). For stimulus decoding, voxels were pooled across the left and right hemisphere. For action decoding, the number of voxels was evenly split between the left and right hemispheres so as to evenly distribute feature-selectivity between classes. We used these voxel selection criteria because they were based on well-established functional properties of retinotopic visual cortex and somatotopic motor cortex, which we verified, separately for each individual, for areas V1 (stimulus) and M1 (action; Figure S9).

We then used an iterative procedure to estimate single-trial response amplitudes (beta weights) for each of the selected voxels, by fitting one GLM for each trial (Mumford et al., 2012). Each of those GLMs contained two regressors: one stick function for the trial of interest, and another stick function for all remaining trials, whereby both regressor time series were convolved with the individual HRF estimate (see *Individual HRF estimation* above). Single-trial regression was performed on all runs (including both the instructed and inferred rule runs) concatenated within sessions, but not across sessions, under the assumption that the discriminant pattern may differ between sessions. The resulting trials x voxels matrix served as the data for classification.

Decoding was cross-validated with ten folds. The conditions in the training data, but not the testing data, were balanced by up-sampling the condition with fewer trials, and the training and testing data were subsequently z-scored separately. Decoding accuracy was then computed in the testing data, as the percentage of correctly classified horizontal choice gratings (or leftward actions) plus the percentage of correctly classified vertical choice (or rightward actions) gratings, divided by two. For each type of classification (stimulus orientation or action), this yielded a participant x session x fold x ROI matrix of classification accuracies, which was then averaged across sessions and cross-validation folds. We assessed its significance for each ROI using permutation testing (10,000 iterations) by comparing it to chance level (50% for both stimulus and action decoding). Similar results in terms of direction and significance were obtained using alternate numbers of included voxels (50, 75, 100, or 200).

#### Removal of feature-specific evoked responses

We again used deconvolution to isolate the ongoing fluctuations of each voxel’s activity in order quantify the correlation structure of these fluctuations (see subsequent sections). We here use the term ‘ongoing activity’ as a shorthand to refer to activity unrelated to the events of the primary choice task (grating stimuli or button presses), acknowledging that these fluctuations may be partly driven by the SR-mapping cue (instructed rule) or the evidence samples for the higher-order decision (inferred rule). Responses to such events are not expected to generate activity patterns encoding one or the other stimulus orientation, or one or the other action. Thus, such responses do not confound our analyses of correlations between stimulus and action decoder outputs described in subsequent sections.

We estimated the mean orientation-specific or action-specific evoked responses and then subtracted these feature-selective evoked responses from the voxel time series. For each type of ROI (visual field maps or action-related), we thus used the procedure described above (see (*Deconvolution of evoked fMRI responses*), but now to construct two separate matrices S, one per feature (horizontal / vertical grating orientation, or left / right response hand). These two matrices were then concatenated horizontally to produce a new composite design matrix X with dimensionality N x 2P. The time series of ongoing activity (i.e., feature-specific evoked responses removed) was then computed as the residual (*ε*) (Dale, 1999):

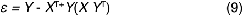

where ^+^ denotes pseudoinverse, ^T^ denotes transposition, and *Y* was the up-sampled, detrended, and z-scored voxel time series concatenated across the complete session. This procedure was repeated for both sessions.

To verify that this procedure effectively removed feature-specific evoked activity (Figure S3), we projected the SVM decision function onto the time series of multi-voxel patterns (voxels used for stimulus and action decoding, see above) and computed the mean decoding accuracy across trials, time-locked to grating onset (visual cortical areas) or button press (action-related ROIs). This was done once for the full voxel time series and once for the residual voxel time computed via eq. 9.

#### Correlation of stimulus- and action-selective voxel groups

To illustrate the rationale behind our feature-specific correlation analyses in a simple format for an example participant and area pair (Figure 2g), we selected the 150 voxels from across early visual cortex (V1-V4) that exhibited the largest responses to horizontal gratings, and another 150 voxels from the same ROI with maximal responses to vertical gratings. Likewise, we selected 150 voxels in left PMd with maximal responses during right button presses, and 150 voxels in right PMd with maximal responses during left button presses. These response amplitudes were estimated with GLM regressing these trial events (grating orientations or actions) on the time series of all voxels within a given ROI, after convolution with the individual HRF estimate. We first removed feature-specific evoked responses (eq. 9) and then the mean activity across all selected voxels (via linear regression). The resulting voxel time series were then selectively averaged for each of the two voxel groups (i.e., across all vertical-preferring and all horizontal-preferring voxels (visual areas) or across all right-preferring and all left-preferring voxels (action-related areas).

We next created a time series that encoded active rule at each given time point in the fMRI data (rule 1 = -1; rule 2 = 1), convolved it with the individual HRFs, and took the sign of this vector to define alternating data segments (mean duration: 23.5 s) corresponding to the two active rules. We correlated the time series of all four pairs of stimulus- and action-selective voxel groups, separately for the two types of data segments corresponding to the two active rules. To this end, we concatenated the segments of each type and then computed correlations per type.

#### Correlation of stimulus and action decoder outputs

Our main analyses computed the correlations between the graded outputs of stimulus and/or action decoders applied to the fluctuations of multi-voxel activity patterns (i.e., after removal of feature-specific evoked responses; see above). The analysis pipeline is illustrated in Figure 2h. For each participant, ROI, and experimental session, we projected the decision functions from the stimulus and action decoders trained on the evoked responses (see *Decoding of stimulus and action from patterns of evoked responses*) onto our estimates of ongoing multi-voxel activity patterns (see *Removal of feature-specific evoked responses*). This yielded a scalar time series that quantified the graded decoder output, which was signed and could be interpreted as the instantaneous tendency of the ongoing population activity towards one or the other stimulus orientation (visual decoders) or one or the other action (action decoders). Given the way we computed the decoders, high values indicated an ongoing activity pattern resembling responses to vertical gratings (left-hand button presses) and low values indicated an ongoing activity pattern resembling responses to horizontal gratings (rightward action).

To assess rule-specific inter-area correlations, we segmented the time series corresponding to the active rule, as instructed to (instructed rule task) or inferred by (inferred rule task) the participant. For the instructed rule task, the segmentation was implemented as described above (*Correlation of stimulus- and action-selective voxel groups*). For the inferred rule task, we replaced the block vector for active rule by the time series of *L* from the fitted behavioral model, applied convolution with the individual HRFs (to account for hemodynamic lag), and took the sign of the resulting time series. We then correlated the continuous decoder outputs between all pairs of areas, separately for the two segment types corresponding to (belief in) rule 1 or rule 2, again after concatenating the segments of each type. For the instructed rule task, segments had a mean duration of 23.5 s, and for the inferred rule task, they had a mean duration of 21 s.

#### Similarity between correlation patterns across visual-action ROI pairs

We used correlation analyses to quantify the similarity between patterns of correlations of decoder outputs across all visual-action ROI pairs, computed for either the intrinsic signal fluctuations after removal of evoked responses (‘intrinsic correlations’), as used in the main analyses, or for trial-evoked responses (‘evoked correlations’). We estimated the evoked responses by projecting the stimulus and action classifiers onto the voxel-level data, without residualization. The resulting decoder output time series were epoched, time-locked to stimulus onset, and separately averaged across trials for all four unique stimulus-rule combinations. For each rule, we then concatenated the resulting average decoder responses for the two stimuli, yielding two time series (one per rule) for each ROI. These time series should correlate positively between visual-action ROI pairs for rule 1, and negatively for rule 2, simply due to the rule-consistent flipping of stimulus-response associations in subjects’ behavior. We finally used Pearson correlation to quantify the spatial similarity of this pattern of evoked correlations (Figure S4) with the pattern of intrinsic correlations (gray outline in Figures 3c and 5c).

To quantify the across-trial consistency of evoked correlation patterns in an analogous fashion, we randomly sampled 50% of trials, averaged graded decoder output across trials and computed the pattern of evoked correlations as described above. This pattern was then correlated with evoked correlation pattern for the other 50% of trials. This was repeated 1,000 times for each per participant, to create a null distribution of expected evoked correlation patterns. The average of the null distribution served as the participant-specific estimate of the across-trial similarity (correlation) of evoked correlation patterns, and was close to 0.7 for both the instructed rule (Figure 3d, green) and inferred rule (Figure 5d, green) tasks.

We used an analogous approach to quantify the similarity of patterns of intrinsic correlations between the instructed and inferred rule tasks. To this end, the group average pattern of stimulus and action decoder output correlations in instructed (gray rectangle in Figure 3c) was correlated with the corresponding pattern from inferred (gray rectangle in Figure 5c), and the significance of this correlation was assessed by comparing it to a null distribution that was generated by shuffling the ROI-pair labels at the single-participant level (10,000 iterations).

#### Time-variant correlation of decoder outputs

To estimate stimulus and action decoder correlations in a time-variant manner, we computed a measure of the correlation of graded decoder outputs at a given time point *t* as follows:

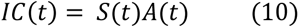

where *S* and *A* were the z-scored time series of the stimulus (*S*) or action (*A*) decoder outputs. Because the time series *IC*(*t*) averages to the Pearson correlation coefficient, it is a measure of the instantaneous coupling between the decoder outputs (Zamani Esfahlani et al., 2020).

#### Prediction of active rule or belief from decoder correlations

We used cross-validated multiple regression to test whether the fluctuations of time-variant correlations between stimulus and action decoders (see eq. 10), and/or of the continuous decoder outputs themselves, predicted the instantaneous belief about the active rule (inferred rule task) or the currently active rule (instructed rule). The fluctuations of decoder correlations contained correlations of 84 pairs of ROIs, which tended to be similar (Figs. 3 and 5). To reduce the dimensionality of the data (i.e., the feature set for the decoder), we submitted the matrix of time-variant correlations of all stimulus-action pairs, estimated with eq. 10, to PCA. We selected the number of components used for prediction by comparing the eigenvalues *λ* to a theoretical noise distribution *ρ* (Mitra and Pesaran, 1999):

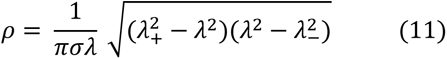

where:

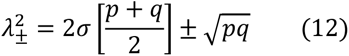

and σ was the standard deviation of *λ*, and *p* and *q* were the dimensions of the covariance matrix of correlated decoder outputs. A scalar-multiplied version of *ρ* was fit to *λ*. The components for which *ρ* < *λ* were taken as ‘signal’ (Mitra and Pesaran, 1999), and their time series were included in the regression model. A procedure where we fixed the number (12) of selected components based on the group-average yielded identical results in terms of direction and significance of effects.

We quantified the contribution of the decoder correlations (*m* selected components, see above), stimulus decoders, and action decoders, to the prediction of the graded belief about the rule (inferred rule) or the currently active rule (instructed rule). For the inferred rule task, we used linear regression:

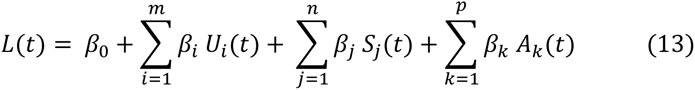

where *L(t)* was the belief time series extracted from the normative model fit for each participant, convolved with the participant-specific HRF (to account for hemodynamic lag), *U* was the set of component time series (quantifying decoder correlations) and *i* indexed the components, *S* was the set of n stimulus decoder outputs from visual cortical areas indexed by *j*, and *A* was the set of *p* action decoder outputs from action-related areas indexed by *k*.

For the instructed rule task, we used logistic regression:

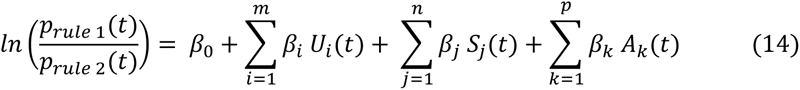

where *prule 1(t)* indicated the probability that rule 1 was active at time point *t* and correspondingly for *prule 2(t),* switching at a mean interval of 23.5 s. The rule time series was convolved with the individual HRF estimate and binarized (see *Correlation of stimulus- and action-selective voxel groups* above) to account for hemodynamic lag.

For both tasks, we used a ten-fold cross-validation procedure to compute prediction accuracy. We divided the data into ten segments of equal length (∼3 minutes each for the inferred rule task, ∼2 minutes for instructed). Each of these segments contained the to-be-predicted variable (HRF-convolved) and the regressor time series (i.e., the selected components as well as the local decoder outputs). For each fold, nine segments were used for training the prediction model, by concatenating the nine segments and computing beta weights (using eqs. 13 or 14). We then used the remaining segment for prediction, by applying the beta weights obtained during training in eqs. 13 and 14. Prediction accuracy was quantified as the mean percentage of correctly predicted rule 1 and rule 2 (chance = 50%; instructed rule), or the correlation between predicted belief and observed belief (inferred rule).

Predictions during testing (in the held-out segment) were either obtained based on all regressors (*U, S, and A*), based on only decoder correlations (*U*), or based on either of the local decoder outputs (*S* or *A*). For each fold, the time series of all regressors, or of *U*, *S*, or *A* individually, that were measured in the testing segment, were multiplied with the beta weights estimated from the training data. The accuracies of these predictions were then calculated, and averaged across the ten folds.

#### Dynamics of rule-specific correlation patterns surrounding changes in rule and belief

We analyzed the dynamics of rule-specific correlation patterns during trials of the primary (orientation discrimination) task that surrounded a change in active rule (instructed) or the sign of *L* (inferred). We focused on two trials before and two trials after rule (belief) change. For each of these trials we computed the correlations between stimulus and action decoders in a time window from -6.8 to +6.8 s from trial onset. This window corresponded to the minimum inter-trial interval and thus prevented overlap between successive trials. We then sorted trials contingent on the change in rule or belief (from rule 1 to rule 2, or vice versa). The timing of these changes in active rule (or sign of *L*) were corrected for the hemodynamic delay. We then computed the average change from trial +1 to +2 post-switch after sign-flipping the values for switches from rule 1 to 2, such that positive values indicated increased coupling from trial +1 to trial +2 (Figure 6h and 6j). These values were then compared to 0 using permutation testing.

#### Simulation of the dynamics of correlated decoder output

We simulated data with known properties for 100 pseudo-participants (similar properties to our participants) to constrain the interpretation of the observed dynamics or rule-specific correlation patterns, mimicking the experimental setting of the instructed rule task. We defined a trial sequence with randomly chosen stimuli in which the active rule alternated every two trials.

Stimulus timing, run duration and the number of sessions per pseudo-participants was identical to the actual data in the instructed rule task. SR-mapping cues appeared in between mini-blocks to signal the rule change. We selected an accuracy level from between 95% and 100% on a given session and inserted error trials in random locations to match this accuracy level.

Neural population activity was simulated for two regions each consisting of 10 voxels, one stimulus-coding and one action-coding region, referred to as ‘visual cortex’ and ‘motor cortex’ in the following. The neural data consisted of 4 components: 1) a response to stimulus onset (visual cortex) or action (motor cortex) where each voxel had a preferred stimulus (or action) that determined its response amplitude. The pattern of stimulus (or action) preferences across voxels constituted the to-be-decoded pattern. 2) A response to the SR-mapping cue of positive uniform random amplitude. 3) Intrinsic fluctuations of the stimulus (visual cortex) or action (motor cortex) encoding pattern: these fluctuations were a vector of IID noise that was multiplied with the feature-coding pattern. Fluctuations in the action pattern were defined as a noise-corrupted version of the fluctuations in the stimulus pattern such that the two were weakly correlated. We flipped the sign of the intrinsic fluctuations in the action-encoding pattern at the time points when rule 2 was active, which ensured an instantaneous flip in correlation between stimulus and action encoding patterns following a change in active rule. 4) Random IID noise to simulate intrinsic fluctuations other than those that encoded the feature of interest. The amplitude of this noise was set such that post-hoc classification accuracy of stimulus orientation or action was similar to our actual data. For each pseudo-participant and session, we then defined an HRF with randomly chosen parameters: a delay of the peak response between 3 and 6 s, delay of the undershoot between 7 and 15 s, and ratio of the peak response to undershoot between 3 and 8. The neural data were then convolved with the HRF, and down-sampled to the same sample-rate as the actual fMRI data.

We analyzed this synthetic data exactly as the actual fMRI data, including HRF estimation with deconvolution, the removal of feature-specific evoked responses, decoder training and correlation of stimulus and action decoder outputs, and rule change-locked analysis of correlated decoder output.

#### Analysis of errors

Analyses of the relationship between decoder correlations and behavioral errors were only possible for the inferred rule task, not the instructed rule task, because participants only made a considerable number of errors in the former. Because there were more correct trials than error trials, we sought to equalize the contribution of both trial types to the decoders. We thus selected voxels for decoder training based on average regression coefficients in which error-trials were up-weighted to match correct trials. Moreover, decoders were trained using data of all error trials, and a randomly sub-sampled number of correct trials such that both trial types contributed equally to the decoder. This sub-sampling was repeated 1000 times, and the decoder outputs were averaged across sub-sampling iterations.

We computed the correlations between stimulus and action decoders separately for time windows from -6.8 to +6.8 s from the trials of the lower-order task (orientation judgments). This window corresponded to the minimum inter-trial interval and thus prevented overlap between successive trials.

We then sorted the trials based on choice correctness (i.e. consistency of the choice with the active rule). There were between 25 and 74 (mean: 46; SD: 15) error trials per participant. Again, because there were fewer error trials, we computed decoder correlations for correct trials on a randomly sampled subset of trials that was equal in number to error trials. We repeated this procedure 1000 times and then averaged across iterations. Finally, we compared the decoder correlations for each rule, as well as the differences between rules, between correct and error trials.

The relationship between correlated decoder output and the model decision noise parameter was assessed using a permutation-based correlation procedure. We selected time windows surrounding choices (see above), and computed the difference in decoder correlations for the two belief states (favoring rule 1 or rule 2), separately for each participant. This difference score was then correlated in turn with the individual noise parameters (*V*) estimated from the behavioral model fits using Spearman’s rank correlation. We used Spearman’s correlation in this case to down-weight one participant with a comparatively high value of decision noise. Qualitatively identical results were obtained with using Pearson correlation.

In all other cases, we use Pearson’s correlation, because it is the standard in the analysis if inter-regional correlation of fMRI signals.

#### Decoding of rule or belief from local activity patterns

We used support vector classification (instructed rule) or support vector regression (inferred rule runs) to test if the local patterns of fMRI activity encoded the (belief about) active rule. These analyses were run for each ROI of the HCP-MMP1.0 parcellation (Glasser et al., 2016), thus covering the complete cortex. Decoding was performed in two different ways: 1) based on the evoked response to the cue that instructed the participant of the active rule (instructed rule runs), and 2) the continuous fMRI time series, non-residualized for evoked responses (both the instructed and inferred rule runs). All voxels from each ROI were included. We used the same ten-fold cross-validation procedure as used in other analyses (see sections: *Decoding of stimulus and action from patterns of evoked responses*, and *Prediction of active rule or belief from decoder correlations*). The classification accuracy of active rule in the held-out runs (instructed rule runs) and the correlation between SVR-predicted and model-estimated belief in the held-out runs (inferred rule task) was compared to chance (permutation tests, see below) and corrected for multiple comparisons using the false discovery rate (FDR; q = 0.05).

To characterize the coarse spatial distribution of rule (and belief) encoding, we grouped cortical parcels into a set of clusters. These clusters were based on macroscale regions outlined in Figure 1 of the supplementary neuroanatomical results in Glasser et al. (2016). Specifically, low level visual cortex (clusters 1 and 2); higher order visual cortex (clusters 3-5); somatomotor cortex (clusters 6-8), and other areas (all other clusters).

#### Correlation between rule / belief decoder output and stimulus-action decoder covariation

We correlated the continuous output of the rule decoder (instructed) or belief decoder (inferred), computed as above (section *Decoding of rule or belief from local activity patterns*), with the time-variant stimulus-action decoder coupling (computed via eq. 10). This was done for each stimulus-action ROI pair, and each ROI that showed significant prediction scores of rule or belief (orange areas in Figure 8c). For bilateral ROIs (V1 and 6d) we first averaged the belief decoder output across the two hemispheres. Prior to computing the belief decoder output, we removed evoked responses due to the SR-cue in the instructed rule task. We averaged the correlation between belief / rule decoder output, and time-variant stimulus-action decoder output, across all stimulus-action ROI pairs to obtain a final correlation score. This was compared to chance using permutation testing (10,000 iterations).

*Whole brain GLMs*

We fitted two separate voxel-wise GLMs to compute statistical maps of the covariation of fMRI activity with a number of external trial variables (grating stimulus orientations, left and right button presses) or hidden computational variables extracted from the fitted behavioral model. Both GLMs contained four regressors. The first GLM contained one regressor for each unique stimulus-action combination. The second GLM contained the magnitude of belief (|*L*|) and of sensory evidence (|*LLR*|), change point probability (*CPP*), and all grating onsets (stick function). All regressors were convolved with the individual HRFs.

These GLMs were fit separately for each run, and the resulting beta weights were averaged across runs and sessions. Contrasts were tested using FSL’s ‘randomise’ (10,000 iterations) and corrected for multiple comparisons using threshold-free cluster enhancement (Smith and Nichols, 2009).

#### Standard functional connectivity analysis

For a conventional functional connectivity analysis, we computed inter-regional correlations between the voxel-averaged ongoing activity of all ROIs, separately per active rule (instructed rule) or belief-favored rule (inferred rule). We then collapsed across the instructed and inferred rule runs and compared the correlations visual and action-related areas between rules, as we did for the correlations of decoder outputs from the same areas.

### Statistical tests and error bars

#### Within-participant error bars

For within-participant comparisons, we discarded between-participant variance by mean-centering each participants’ data prior to calculating the SEM (Cousineau, 2005). The purpose of this procedure was to ensure that the error bars depict only within-participant, across-condition, variance that was relevant for the statistical comparison.

#### Statistical tests

We used non-parametric permutation tests (Tibshirani and Efron, 1993) with 10,000 permutations for all statistical comparisons, unless mentioned otherwise. Specifically, single-participant correlations were compared with zero or between rules. The significance of decoding and prediction accuracies was tested by comparing the obtained percentage correct to 50% chance level (instructed rule) or the correlation coefficients between predicted and actual beliefs to zero (inferred rule).

For testing the across-participant correlation between decision noise *V* and decoder correlation difference, we repeated the Spearman correlation 10,000 times, but with shuffled *V* values, yielding a null distribution of correlation coefficients. A *p*-value for the observed correlation was finally obtained by comparing that value to the null distribution.

Unless stated otherwise, in all cases where we conducted multiple statistical tests, we applied the FDR correction for multiple comparisons.

## Supplemental Information

**Figure S1.**
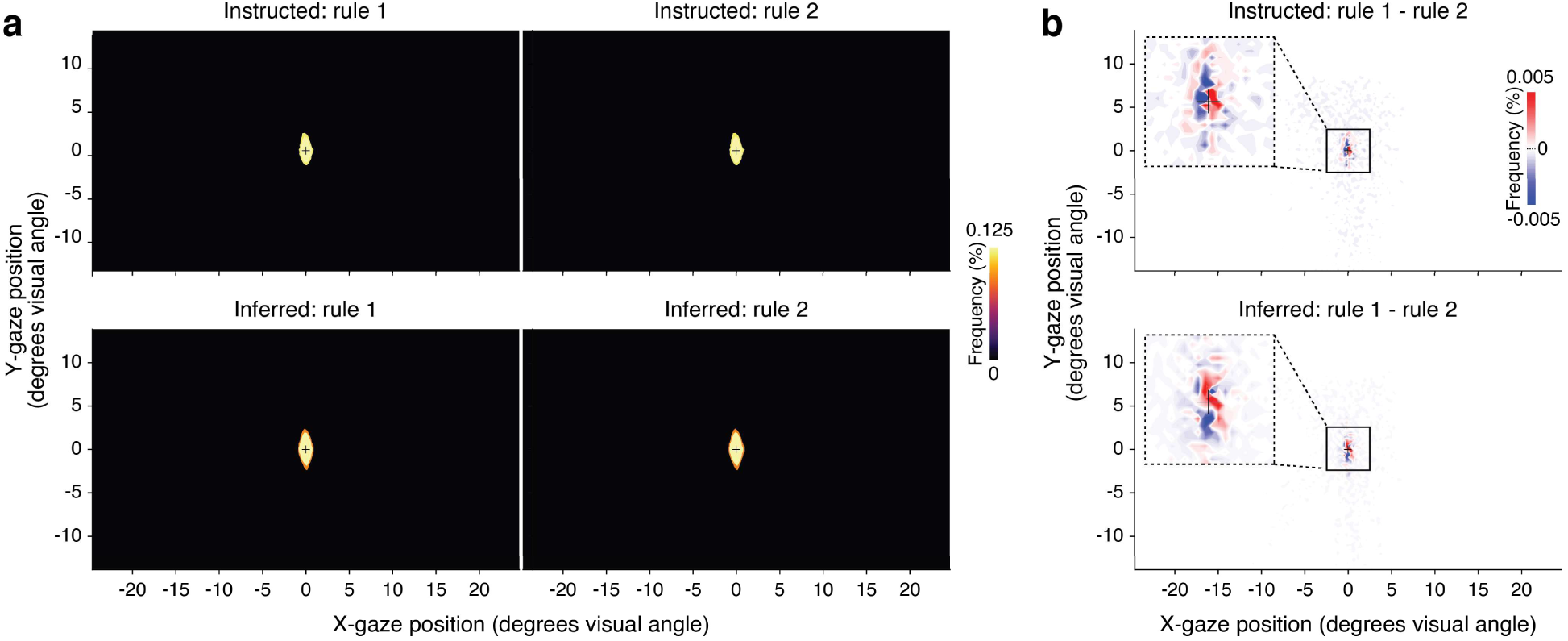
No systematic difference in gaze behavior between rules. **a.** Two-dimensional histograms of eye position across the entire range of the visual display screen. Histograms are shown separately for the instructed rule (top row) and inferred rule (bottom row) tasks, and for rule 1 (left column) and rule 2 (right column). Distributions are centered around the fixation cross, with negligible values beyond 3°, indicating that participants complied with the instruction to maintain central fixation, except for occasional microsaccades. **b.** Distributions of the difference in eye position between rules, separately for the instructed rule (top) and inferred rule (bottom) tasks. Insets in the top left corner of each panel (dashed rectangle) show a close-up of the central part of the difference distribution (solid rectangle). No systematic pattern is evident in the differential distributions. We also found no statistically significant (*p* < 0.05, FDR corrected) difference in frequency between rules for any eye position bin.

**Figure S2.**
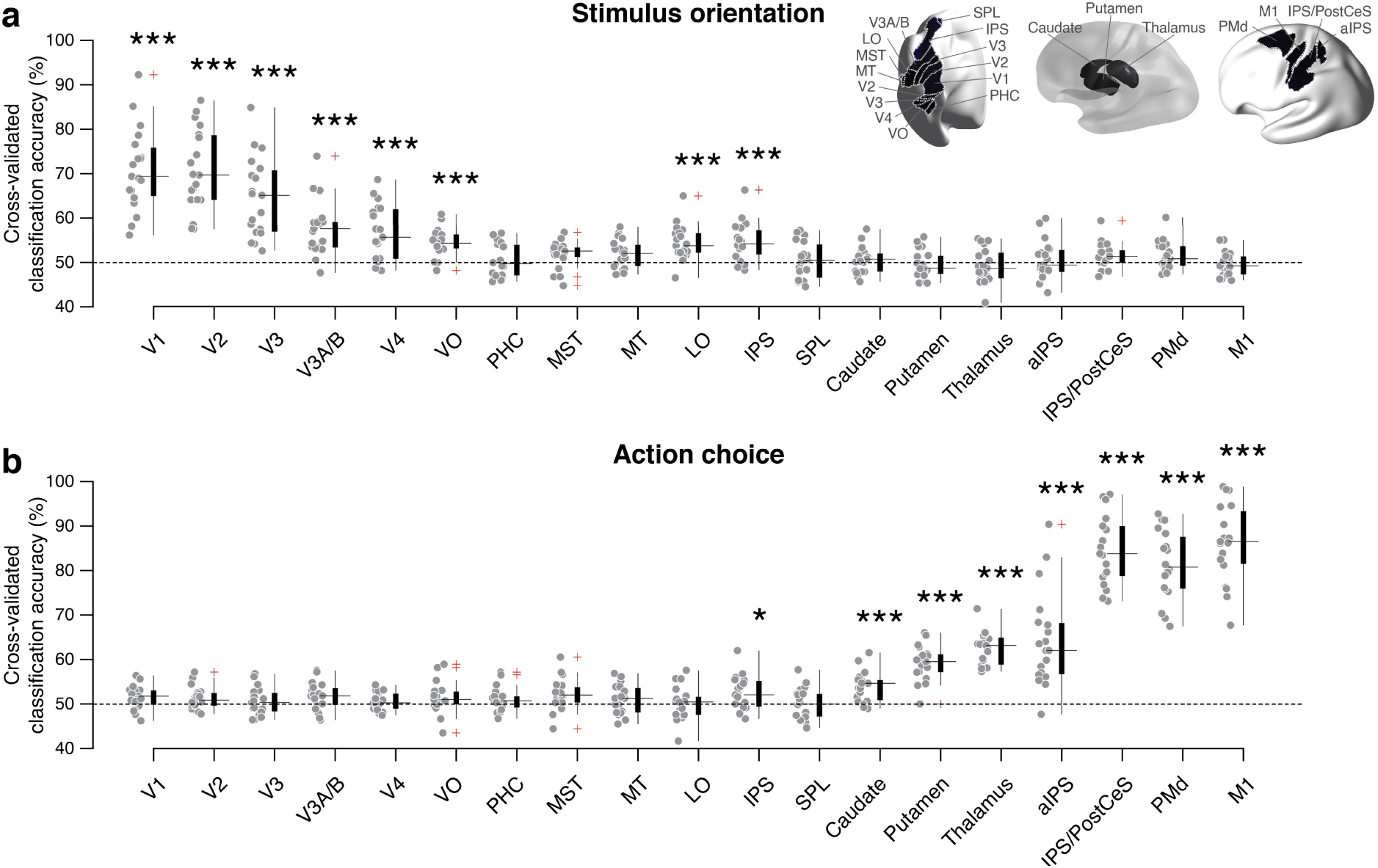
Cross-validated classification accuracy of a. stimulus orientation, and b. action choice. As expected, classification performance for stimulus orientation was highest for early visual cortex, decreased along the visual cortical hierarchy and was at chance in action-related regions. Conversely, classification performance for action choice increased along the visuo-motor pathway, and peaked in primary motor cortex. Gray dots show individual participants. The whiskers extend to the most extreme values non-outlier values of each distribution. Outliers are shown in red. Asterisks show FDR-corrected *p*-values: \**p* < *0.05*, \*\**p* < 0.01, \*\*\**p* < 0.001

**Figure S3.**
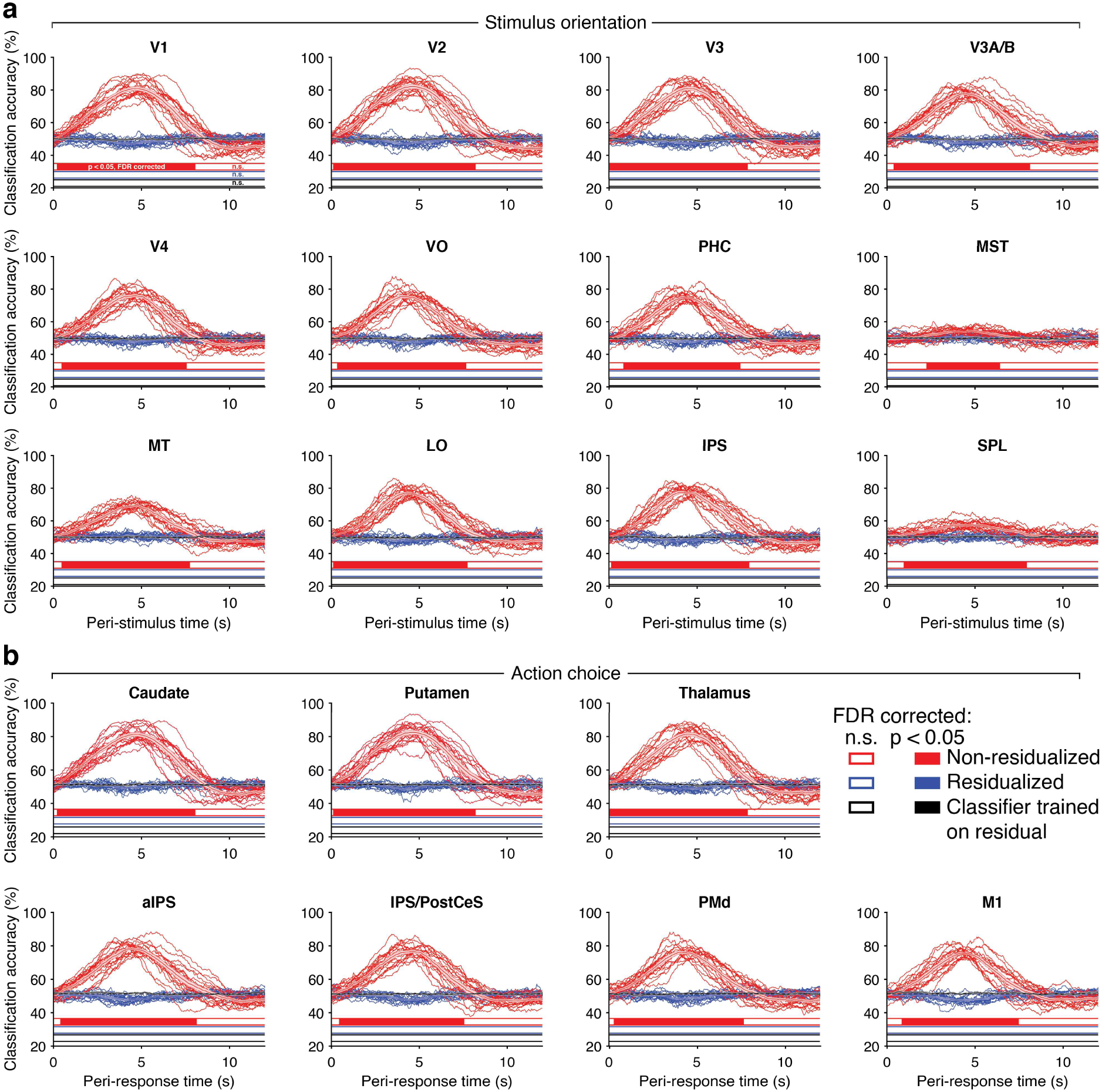
Effective removal of feature-specific evoked responses. **a.** Mean stimulus orientation decoding performance time-locked to grating onset for visual field maps. **b.** Mean action decoding performance time-locked to button presses for action-related ROIs. Red, evoked decoding performance computed based on full voxel time series. Blue, decoding performance of the classifiers trained on non-residualized data, tested on residualized data (i.e., time series after removal of evoked responses). At no time point did decoding performance for residuals deviate significantly from chance in any ROI. Black, as blue, but showing decoding performance of classifiers trained and tested on residualized data (only group average). Again, at no time point did residual decoding performance deviate from chance in any ROI. Thin lines, individual participants; shaded areas, SEM. Together, the results indicate that our analysis effectively removed any feature-specific evoked responses (i.e., discriminant patterns systematically time-locked to stimulus onsets or button presses), regardless of whether the classifiers were trained on the non-residualized or residualized data. This validates the residual activity time courses used for correlation analyses as estimates of ongoing fluctuations in activity (i.e., activity unrelated to the trial events).

**Figure S4.**
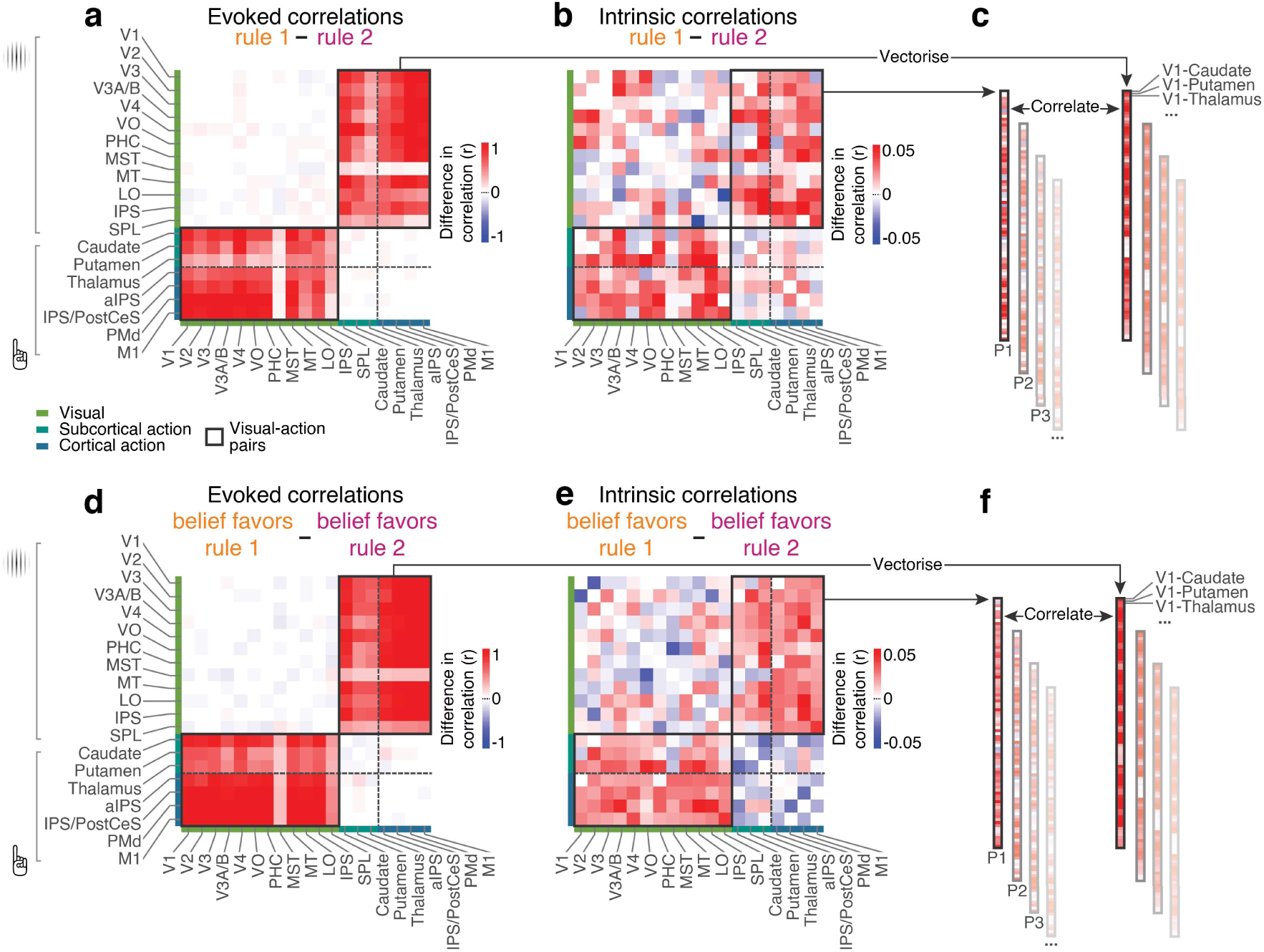
Rationale of control analysis of evoked responses. **a.** Evoked correlation pattern (decoder outputs for evoked responses) in the instructed rule task. **b.** Intrinsic correlations (decoder outputs for intrinsic signal fluctuations) in the instructed rule task (data from Figure 3c; see Main Text), for comparison with panel a. **c.** The edges in panels a and b that link the stimulus and action regions were vectorized, and correlated with one another to quantify their spatial similarity. This yielded a correlation coefficient for each participant (Px), that was compared to zero. **d.** Same as panel a, but for the inferred rule task, and split by dominant belief rather than active rule. **e.** Data from Figure 5c (see Main Text), for comparison with panel d. **f.** Same as panel c, but for the inferred rule task, and split by dominant belief rather than active rule.

**Figure S5.**
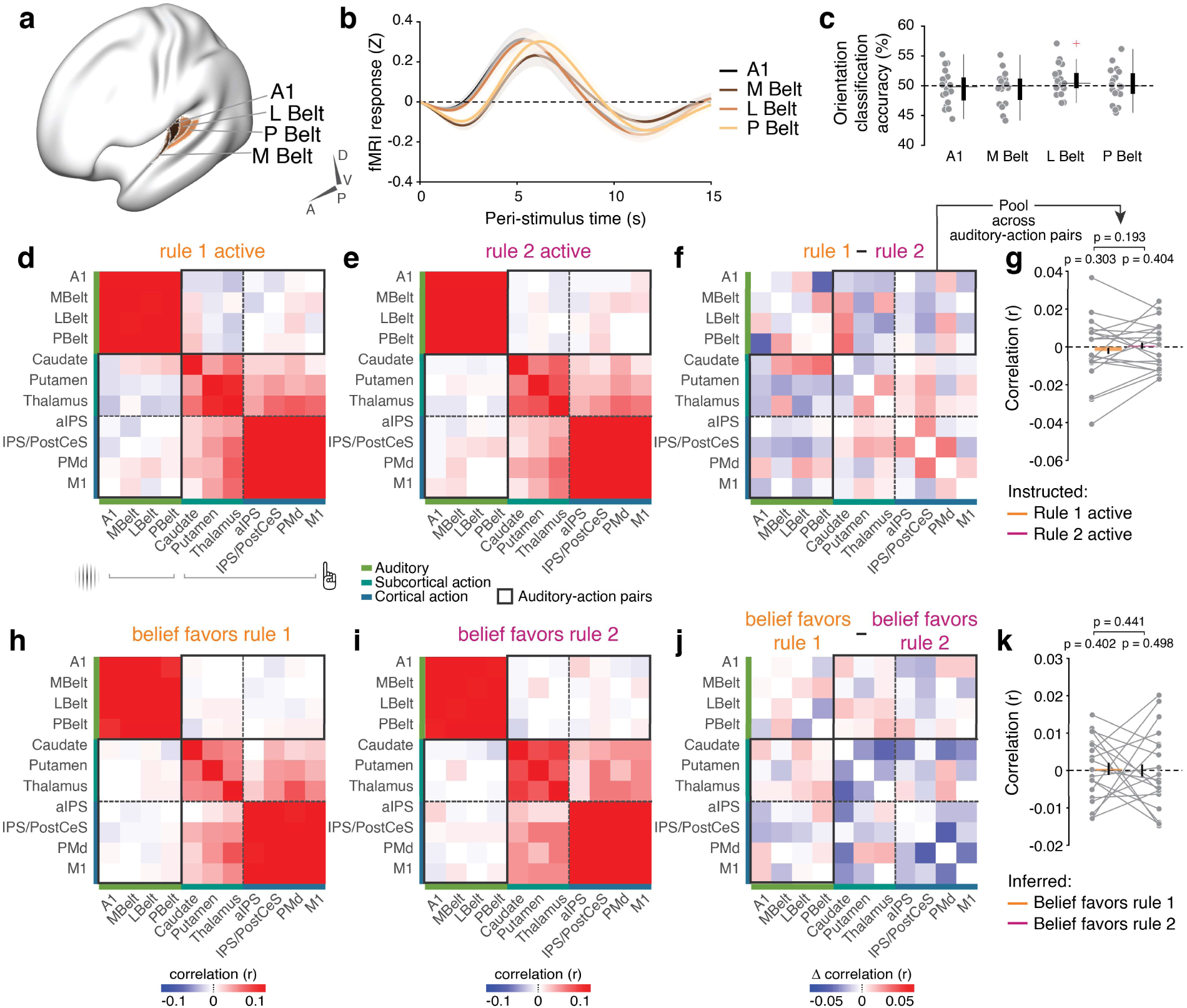
Control analysis using auditory cortex instead of visual cortex. **a.** Regions of interest in auditory cortex that were used to replace visual cortical regions, and entered into stimulus-classification analysis. **b.** Stimulus-locked responses in auditory regions obtained with deconvolution. Auditory regions showed robust responses following trial onset. Error bars, SEM. **c.** Cross-validated stimulus orientation decoder performance. The responses in auditory cortices to stimulus onset were uninformative about stimulus orientation (as expected). Gray dots show individual participants. The whiskers extend to the most extreme non-outlier values of each distribution. Outliers are shown in red. **d.** Correlation matrix of graded decoder output for stimulus and action in the instructed rule task when rule 1 was active. **e.** Same as panel b, but for rule 2. **f.** The difference in coupling between rule 1 and rule 2. The gray outline delineates the edges of the matrix that link stimulus and action ROIs. **g.** Correlation of the graded outputs of a stimulus orientation decoder (trained on auditory cortex) and action decoder. Thin lines, individual participants. Bars, group mean. Error bars, within-participant SEM. **h.** Correlation matrix of graded decoder output for stimulus and action in the inferred rule task when model-derived belief favored rule 1. **i.** Same as panel b, but for rule 2. **j.** The difference in coupling between rule 1 and rule 2. The gray outline delineates the edges of the matrix that link stimulus and action ROIs. **k.** Correlation of the graded outputs of a stimulus orientation decoder (trained on auditory cortex) and action decoder. Thin lines, individual participants. Bars show, mean. Error bars, within-participant SEM.

**Figure S6.**
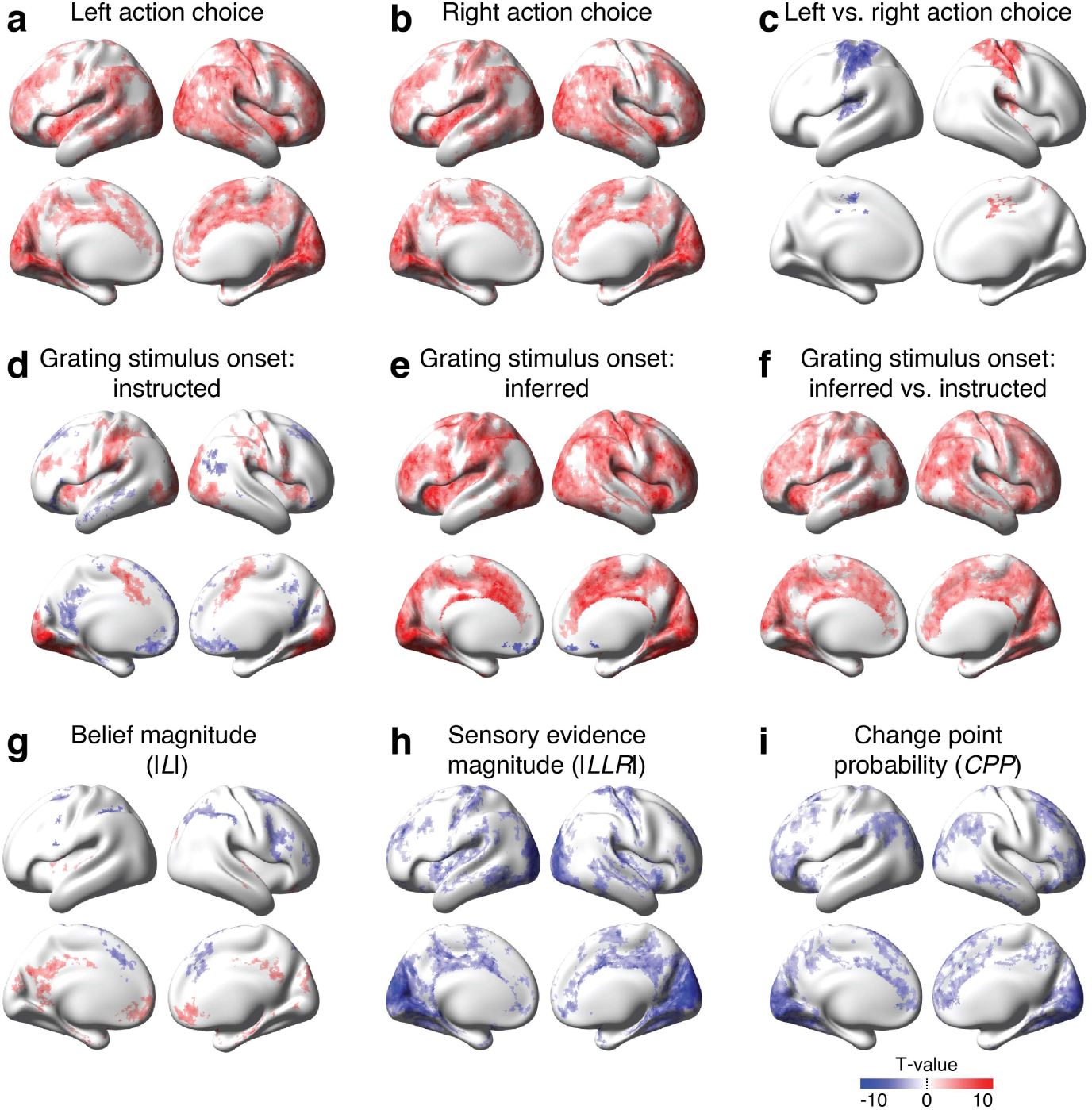
Voxel-level regression analysis of fMRI responses evoked by action, stimulus onset, and computational variables. **a.** Left hand button presses. **b.** Right hand button presses. **c.** The difference between left- and right-hand button presses, showing the expected hand movement-selective lateralization of activity within sensory-motor cortical areas including the action-selective regions of interest used here (PMd, IPS/PostCes, aIPS). **d.** Grating stimulus onset in the instructed rule task. **e.** Grating stimulus onset in the inferred rule task. **f.** Difference between stimulus-locked responses from the instructed and inferred rule tasks. This widespread differential activity likely reflects the increased attentional demand in the inferred rule task. **g.** Magnitude of model-derived belief, showing positive effects in posterior cingulate and ventromedial prefrontal cortices and negative effects in posterior parietal and lateral prefrontal cortices. This cortical distribution resembles the sign-flipped map of effects previously shown for relative uncertainty in a similar learning task (McGuire et al., 2014). This can be understood by noting that belief magnitude in our task roughly corresponds to the complement of relative uncertainty (Murphy et al., 2021). **h.** The magnitude of the sensory evidence in the form of a log-likelihood ratio between rules. **i.** Change point probability, the posterior probability that a change in rule has just occurred given the sensory evidence and belief. Panels a-f were computed in one regression model. Panels g-i were computed in another regression model. All colored regions indicate significant co-variation with local fMRI responses, corrected for multiple comparisons using threshold-free cluster enhancement.

**Figure S7.**
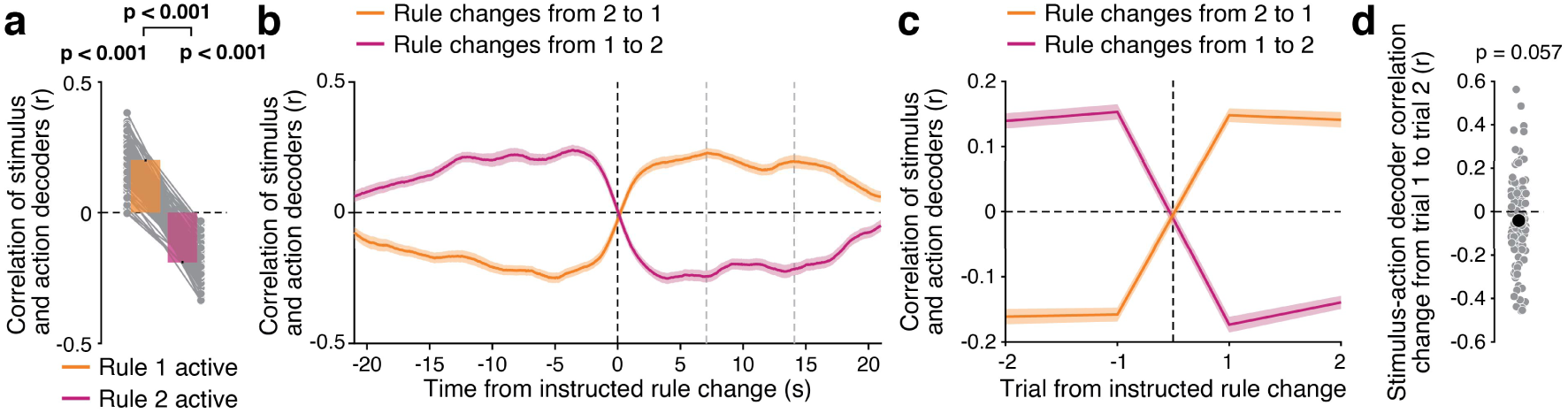
Simulation of rule-specific correlations in synthetic data (see Methods). **a.** Correlation of stimulus- and action decoder outputs for the two rules. Same format as for the data shown in the main figures (e.g., Figure 2i). Gray lines, individual pseudo-participants; bars, group average; error bars, within-participant SEM. **b.** Average time-resolved correlation of decoder outputs locked to the presentation of the SR-cue (HRF-delay corrected). Gray vertical dashed lines indicate the median times of the two trials following the change in rule. The simulated underlying neural correlations flip their sign instantaneously at the time of rule switch and are then sustained at a fixed level until the next switch (not shown). Due to convolution with the HRF, the strength of the correlation increases (or decreases) gradually but reaches a maximum well before the onset of the first trial after rule switch. **c.** Stimulus-action decoder correlation as function of trial from rule changes in the instructed rule task. (c.f. Figure 6g). c-d: Shaded areas, between-participant error bars. **d.** Change in correlation from trial 1 to trial 2 after instructed rule change (c.f. Figure 6h). There is no significant change in decoder output correlation from one trial to the next after rule change, with a trend towards a decrease, in sharp contrast to the data (Figure 6g-j). This indicates that the findings from Figure 6g-j cannot be accounted for by sluggish and accumulating hemodynamics. Gray dots, individual pseudo-participants; Black dot, group average.

**Figure S8.**
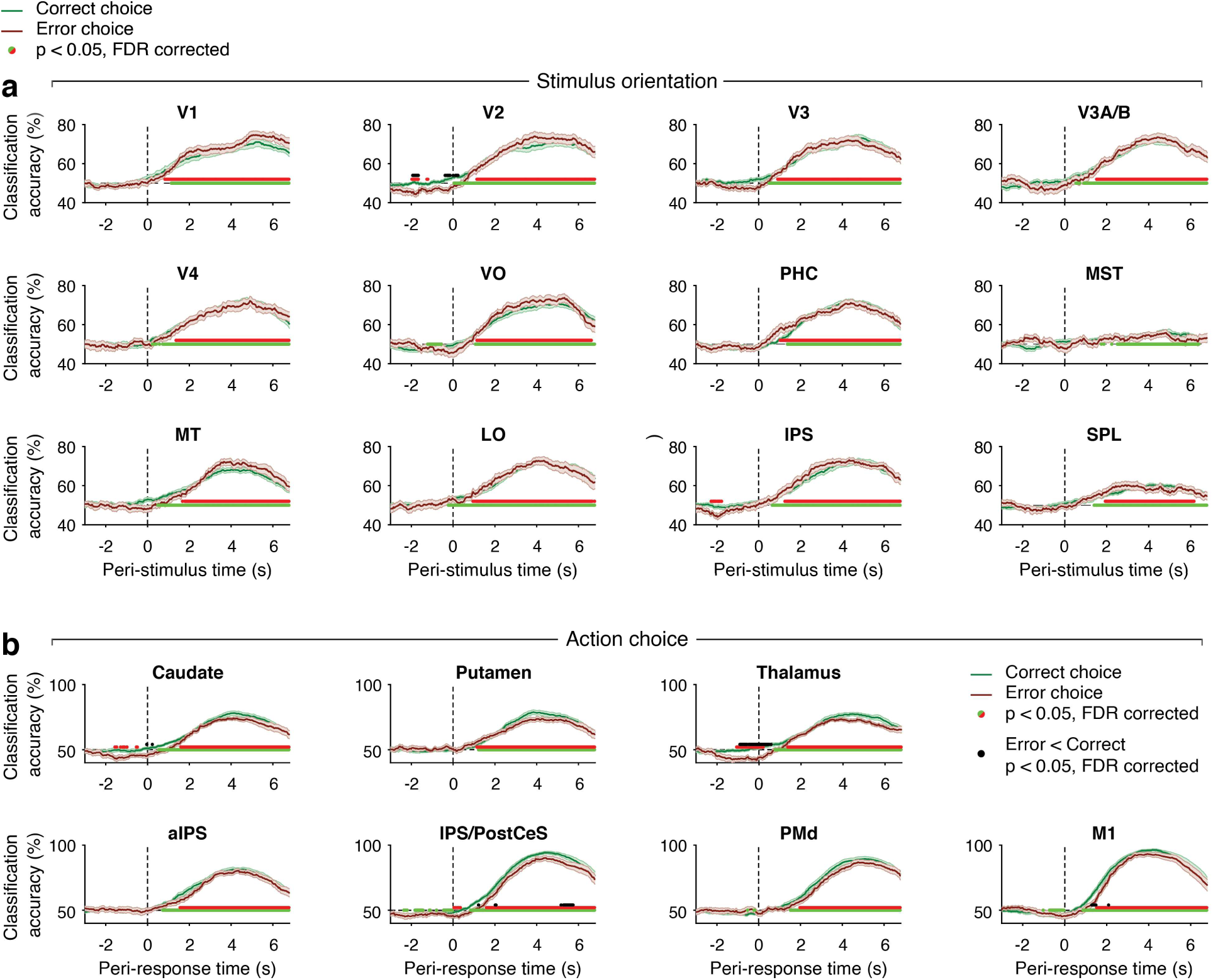
Classification accuracy on correct and error trials in the inferred rule task. **a.** Cross-validated classification accuracy for stimulus orientation in visual cortical areas. Stimulus decoding was significant on, and did not differ between, correct and error trials for early visual cortex (V1-V4). In higher-tier visual cortical area, stimulus decoding was generally weaker, but nowhere was there a significant difference between correct and error trials. **b.** Cross-validated classification accuracy for action in action-related brain regions. Action decoding was significant for both trial categories in all regions. Yet, action decoding was significantly weaker on error than correct trials within Thalamus, IPS/PostCeS, and M1. Overall, this pattern of decoding results indicates that noise corrupting the sensory representation of stimulus orientation judgement was a negligible factor limiting behavioral performance of the inferred rule task, in line with the near-perfect performance based on the same stimuli on the instructed rule task. At the same time, erroneously chosen actions were encoded less precisely in action-related cortical regions, similar to effects observed for high vs. low decision confidence in motor cortex (Wilming et al., 2020). Error bars, within-participant SEM.

**Figure S9.**
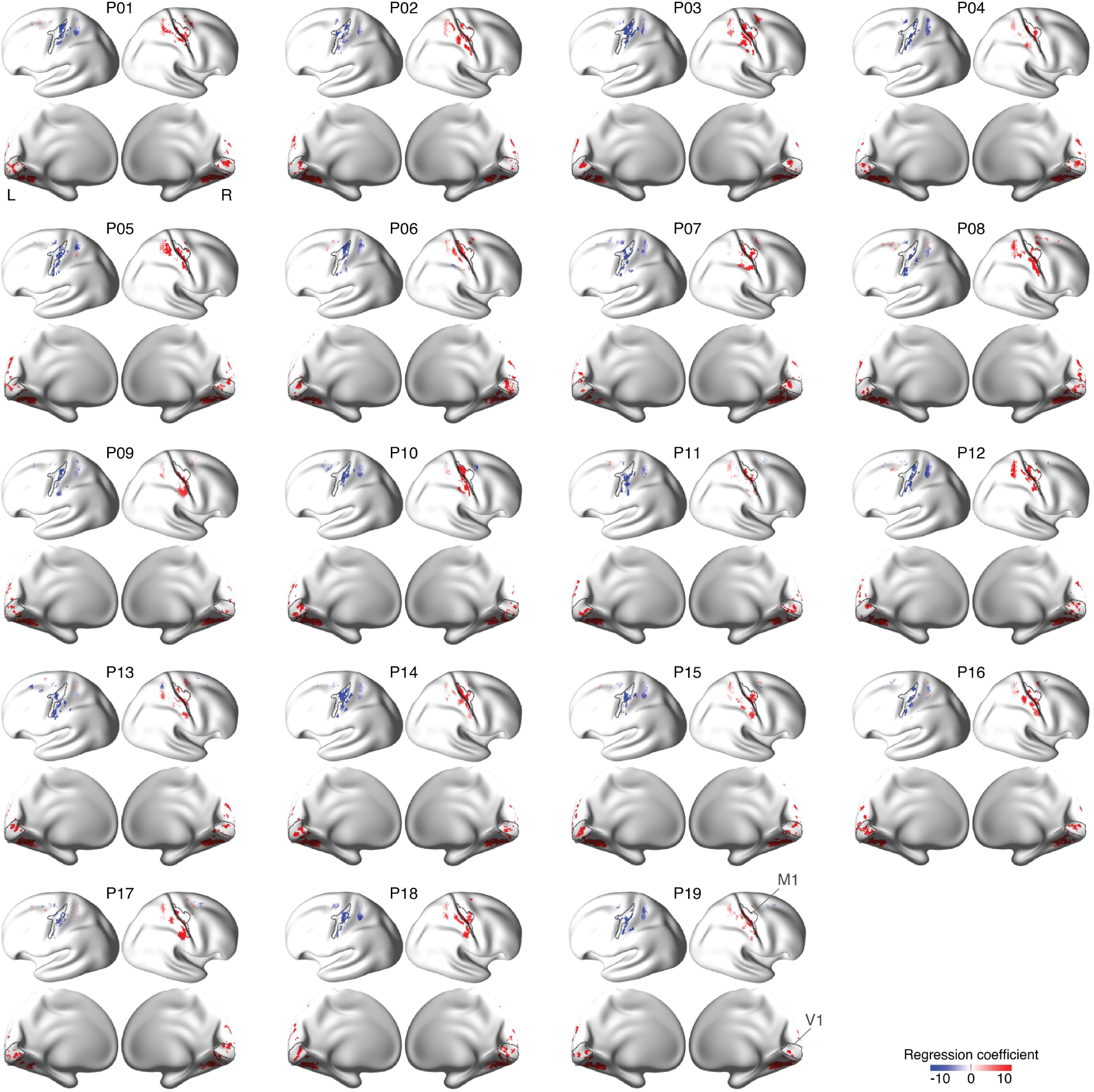
Maps of the voxels selected for decoding for each participant. For each participant, the top row shows the contrast of left vs. right button press (with M1 outlined) and the bottom row shows the response to the onset of the grating stimulus (V1 outlined). Both features of evoked fMRI responses are as expected for each individual included in this study. These results verify the biological plausibility of our voxel selection based on well-characterized properties of evoked cortical responses.

**Table S1.** ROI definition.

| Figure | Label | Test | Tail | *p*-value | Cohen's *d* | 95% CI |
| --- | --- | --- | --- | --- | --- | --- |
| 2d | Rule 1 vs rule 2 | Permutation | Two | 0.741 | -0.075124 | -0.9575-0.63176 |
| 2e | Rule 1 vs rule 2 | Permutation | Two | 0.877 | -0.034624 | -0.014404-0.012257 |
| 2i | Rule 1 vs rule 2 | Permutation | One | 0.023 | 0.47334 | 0.0041928-0.051371 |
| 3d | Evoked vs intrinsic | Permutation | Two | 0.560 | 0.12965 | -0.040756-0.084687 |
| 3d | First vs second half | Permutation | Two | <0.001 | 8.4315 | 0.49529-1.0791 |
| 3d | difference | Permutation | Two | <0.001 | -3.5606 | -1.0753--0.45588 |
| 3e (left) | Rule 1 | Permutation | One | 0.006 | 0.6141 | 0.0025971-0.013267 |
| 3e (left) | Rule 2 | Permutation | One | <0.001 | -1.1924 | -0.018767--0.0067626 |
| 3e (left) | Rule 1 vs rule 2 | Permutation | One | <0.001 | 1.2077 | 0.010594-0.03066 |
| 3e (middle) | Rule 1 | Permutation | Two | <0.001 | 3.8662 | 0.042686-0.093187 |
| 3e (middle) | Rule 2 | Permutation | Two | <0.001 | 2.7499 | 0.03987-0.089326 |
| 3e (middle) | Rule 1 vs rule 2 | Permutation | Two | 0.321 | 0.23399 | -0.002255-0.009223 |
| 3e (right) | Rule 1 | Permutation | Two | <0.001 | 2.705 | 0.054426-0.12187 |
| 3e (right) | Rule 2 | Permutation | Two | <0.001 | 2.3963 | 0.050946-0.11645 |
| 3e (right) | Rule 1 vs rule 2 | Permutation | Two | 0.502 | 0.15744 | -0.005504-0.013617 |
| 3f | Rule 1 | Permutation | One | 0.011 | 0.55745 | 0.0028038-0.017159 |
| 3f | Rule 2 | Permutation | One | <0.001 | -0.77601 | -0.021257--0.005729 |
| 3f | Rule 1 vs rule 2 | Permutation | One | <0.001 | 1.2003 | 0.011919-0.03479 |
| 4e | Perfect vs average | Permutation | Two | <0.001 | 3.4758 | 14.0554-33.2212 |
| 4e | Last sample vs average | Permutation | Two | <0.001 | 5.3166 | 12.4909-28.6638 |
| 5d | Evoked vs intrinsic | Permutation | Two | 0.716 | -0.077789 | -0.060319-0.039052 |
| 5d | First vs second half | Permutation | Two | <0.001 | 3.2144 | 0.37749-0.83617 |
| 5d | difference | Permutation | Two | <0.001 | -2.9153 | -0.88174--0.35197 |
| 5e (left) | Rule 1 | Permutation | One | <0.001 | 0.67003 | 0.0032947-0.014098 |
| 5e (left) | Rule 2 | Permutation | One | <0.001 | -1.1957 | -0.016552--0.006147 |
| 5e (left) | Rule 1 vs rule 2 | Permutation | One | 0.002 | 1.1801 | 0.0099246-0.030193 |
| 5e (middle) | Rule 1 | Permutation | Two | <0.001 | 3.2304 | 0.042469-0.094079 |
| 5e (middle) | Rule 2 | Permutation | Two | <0.001 | 3.9349 | 0.042704-0.093741 |
| 5e (middle) | Rule 1 vs rule 2 | Permutation | Two | 0.908 | -0.025933 | -0.0046983-0.003984 |
| 5e (right) | Rule 1 | Permutation | Two | <0.001 | 2.9657 | 0.05113-0.11221 |
| 5e (right) | Rule 2 | Permutation | Two | <0.001 | 3.0817 | 0.056664-0.12575 |
| 5e (right) | Rule 1 vs rule 2 | Permutation | Two | 0.031 | -0.55756 | -0.017333--0.0020876 |
| 5f | Rule 1 | Permutation | One | 0.013 | 0.5043 | 0.0018525-0.014064 |
| 5f | Rule 2 | Permutation | One | 0.001 | -0.74354 | -0.016268--0.0043726 |
| 5f | Rule 1 vs rule 2 | Permutation | One | <0.001 | 0.83184 | 0.0074151-0.029164 |
| 6a | n/a | Permutation | Two | <0.001 | n/a | 0.2354-0.5974 |
| 6c | n/a | Permutation | Two | <0.001 | 0.91712 | 0.017513-0.054644 |
| 6d | Visual | Permutation | Two | 0.670 | 0.090652 | -0.0090066-0.015404 |
| 6d | Action | Permutation | Two | 0.220 | -0.27362 | -0.030854-0.0039692 |
| 6e | n/a | Permutation | Two | <0.001 | 1.0411 | 1.0086-2.9948 |
| 6f | Visual | Permutation | Two | 0.115 | 0.35107 | -0.042664-1.9264 |
| 6f | Action | Permutation | Two | 0.074 | 0.40531 | 0.062856-2.2688 |
| 6h | n/a | Permutation | Two | <0.001 | 1.0243 | 0.0090961-0.02924 |
| 6j | n/a | Permutation | Two | 0.005 | 0.77 | 0.0032925-0.014296 |
| 7c (left) | Rule 1 | Permutation | Two | <0.001 | 0.86487 | 0.013758-0.042126 |
| 7c (left) | Rule 2 | Permutation | Two | <0.001 | -0.70556 | -0.023765--0.0063366 |
| 7c (left) | Rule 1 vs rule 2 | Permutation | Two | <0.001 | 0.84568 | 0.018626-0.06679 |
| 7c (middle) | Rule 1 | Permutation | Two | 0.003 | 0.64135 | 0.0030537-0.012997 |
| 7c (middle) | Rule 2 | Permutation | Two | 0.084 | 0.4188 | 0.000528-0.011432 |
| 7c (middle) | Rule 1 vs rule 2 | Permutation | Two | 0.201 | 0.23044 | -0.0013546-0.0052119 |
| 7c (right) | Rule 1 | Permutation | Two | <0.001 | 0.91088 | 0.0098797-0.029988 |
| 7c (right) | Rule 2 | Permutation | Two | <0.001 | -0.73474 | -0.032626--0.0093453 |
| 7c (right) | Rule 1 vs rule 2 | Permutation | Two | <0.001 | 0.83644 | 0.017776-0.064116 |
| 7d | n/a | Permutation | One | 0.047 | n/a | -0.8070--0.0124 |
| 8e (top) | Visual (low) vs other | Permutation | Two | <0.001 | 1.1754 | 0.4697-1.3565 |
| 8e(top) | Visual (low) vs somatomotor | Permutation | Two | 0.075 | 0.43142 | 0.023243-0.69902 |
| 8e(top) | Visual (low) vs visual (high) | Permutation | Two | 0.003 | 0.8117 | 0.31136-1.1866 |
| 8e (top) | Visual (high) vs other | Permutation | Two | 0.221 | 0.29434 | -0.046009-0.37714 |
| 8e (top) | Visual (high) vs somatomotor | Permutation | Two | 0.089 | -0.41302 | -0.75003--0.018136 |
| 8e (top) | Somatomotor vs other | Permutation | Two | 0.210 | 0.92285 | 0.23839-0.83929 |
| 8e (bottom) | Visual (low) vs other | Permutation | Two | <0.001 | 1.8652 | 0.070932-0.17909 |
| 8e (bottom) | Visual (low) vs somatomotor | Permutation | Two | <0.001 | 1.8268 | 0.065436-0.16251 |
| 8e (bottom) | Visual (low) vs visual (high) | Permutation | Two | <0.001 | 1.5525 | 0.056186-0.14724 |
| 8e (bottom) | Visual (high) vs other | Permutation | Two | 0.001 | 0.88114 | 0.010552-0.038669 |
| 8e (bottom) | Visual (high) vs somatomotor | Permutation | Two | 0.194 | 0.30584 | -0.0032098-0.026833 |
| 8e (bottom) | Somatomotor vs other | Permutation | Two | 0.014 | 0.61885 | 0.0036579-0.021642 |
| 8g | V1 (instructed) | Permutation | One | 0.051 | 0.3636 | 2.9108*10<sup>-5</sup>-0.0058056 |
| 8g | V1 (inferred) | Permutation | One | 0.012 | 0.5231 | 0.00036839-0.0027402 |
| 8g | V1 (instructed vs inferred) | Permutation | Two | 0.506 | 0.1451 | -0.0016187-0.0040371 |
| 8g | 5L (instructed) | Permutation | One | 0.234 | 0.1617 | -0.0012729-0.0035028 |
| 8g | 5L (inferred) | Permutation | One | <0.001 | 0.7265 | 0.0012669-0.0048225 |
| 8g | 5L (instructed vs inferred) | Permutation | Two | 0.109 | -0.2874 | -0.0048578-0.00057832 |
| 8g | 6d (instructed) | Permutation | One | 0.614 | -0.0635 | -0.0026-0.0017959 |
| 8g | 6d (inferred) | Permutation | One | 0.395 | 0.0566 | -0.0017093-0.0023709 |
| 8g | 6d (instructed vs inferred) | Permutation | Two | 0.648 | -0.0999 | -0.0047201-0.0025871 |
| 8g | 10v (instructed) | Permutation | One | 0.336 | 0.0906 | -0.0019504-0.0035126 |
| 8g | 10v (inferred) | Permutation | One | 0.732 | -0.1345 | -0.0032746-0.0014451 |
| 8g | 10v (instructed vs inferred) | Permutation | Two | 0.575 | 0.1213 | -0.0027493-0.0055297 |
| 9e (right) | Rule 1 | Permutation | Two | <0.001 | 3.9668 | 0.26144-0.57885 |
| 9e (right) | Rule 2 | Permutation | Two | <0.001 | 3.8792 | 0.25447-0.56815 |
| 9e (right) | Rule 1 vs rule 2 | Permutation | Two | 0.155 | 0.346 | -0.001084-0.01932 |

**Coloring:** Black: main result; Blue: control result; Green: decoding result. **Cohen's $d$ :** Measure of effect size. $d > 0.2$ : small effect size; $d > 0.5$ : medium; $d > 0.8$ : large; $d > 1.2$ : very large; $d > 2.00$ : huge. **CI:** confidence interval around mean (for comparisons to chance), or around difference of means (for comparisons between conditions).

